# Constraint Semantics for Multi-level Organisation

**DOI:** 10.64898/2026.02.27.708558

**Authors:** Sahil Imtiyaz

## Abstract

Biological organisation is inherently multi-level: molecular processes, membrane dynamics, cellular geometry and tissue context reciprocally constrain one another, often through boundary-mediated feedback. A recurring theme in theoretical biology is that such organisation is not well captured by models that assume a fixed repertoire of variables and a pre-given state space: what counts as a relevant state description can depend on organisational context and history. The principle of biological relativity further sharpens the same challenge from a different angle, emphasising that no level is causally privileged and that cross-level feedback can close into circular causality. These lines of work motivates for a structural multi-level semantics for modeling the biological pathways.

We introduce a constraint-based semantic framework that distinguishes an evolving organisational scaffold—the admissible multi-level patterns and interfaces—from the pathways that traverse and coordinate them. This separation yields mathematical, loop-level diagnostics for boundary-driven circular causality: it identifies when organisational trajectories induce persistent reparameterisations of local state descriptions, and it classifies cyclic regimes into reversible loops, stable history-dependent loops, and unique (rare) organisational reconfigurations. The framework is accompanied by a systematic crosswalk to mainstream causal, dynamical and computational approaches, clarifying what is gained when interfaces and local–global consistency are treated as semantic, rather than purely parametric, structure.

We demonstrate the approach on a canonical excitable-cell exemplar by modelling a single Hodgkin spike as a cross-level interface loop coupling membrane, molecular and cellular constraints. Without re-deriving Hodgkin–Huxley kinetics, the resulting diagnostics provide an explicit semantics for boundary-mediated feedback and spike-induced history dependence, including when cyclic activity imprints persistent changes in effective excitability. Together, the case study and comparisons position constraint semantics as a practical mathematical layer for multi-level biological organisation: compatible with existing mechanistic models, yet designed to expose circular causal closure and organisation-dependent state descriptions that standard formalisms typically leave implicit.

**AMS subject classifications:** 92C30, 92C46, 92B05, 55U10, 55R10

## 1 Introduction

Biological organisation is intrinsically multi-level: ionic currents and channel kinetics, membrane mechanics and geometry, intracellular regulation, and tissue context jointly constrain one another in ways that are not well captured by strictly separable, single-level descriptions [32, 42, 43]. In the modelling paradigm of theoretical biology, living systems are organised wholes in which structure and function are inseparable, causal relations often operate as *enablements* (opening and reshaping possibilities rather than entailing trajectories), and the relevant observables and admissible behaviours may change with the organisation itself [31–33]. In physiology, Noble’s principle of *biological relativity* makes the methodological demand explicit: there is no privileged causal level, and causal efficacy is distributed across levels through reciprocal constraint propagation and closure into *circular causality* [36, 42, *43]*.

A persistent methodological gap follows. Many widely used formalisms in causal inference, computation, and dynamical systems presuppose some combination of: a fixed variable set, a fixed phase/configuration space, boundaries and interfaces that enter only parametrically, and cycles that are treated indirectly (for example by time-unrolling, equilibrium closure, or embedding feedback into enlarged Markov states) [11, 13, 21, 25, 47, 68]. These assumptions are often appropriate for mechanistic or predictive tasks, but they make it difficult to represent *active interfaces, multi-level constraint closure*, and *organisation-dependent changes* in what counts as a state—precisely the phenomena emphasised in theoretical biology and in Noble’s critique of single-level causation [32, 36, 43]. Tables 2 and 3 (Section 2) consolidate these recurring assumptions across graphical/structural causality [47, 56], time-series and dynamical-systems causality [13, 30, 60], information-theoretic and multi-scale approaches [17, 18, 55], concurrency/computation [48, 64, 66], and (for completeness) operational quantum-causal formalisms [7, 29, 46]. Across these very broad span of successful formalisms, a shared assumption is prestated variables/state space + interfaces treated parametrically + cycles handled indirectly — which does not address with the challenges put forth in theoretical biology.

**Table 1:**
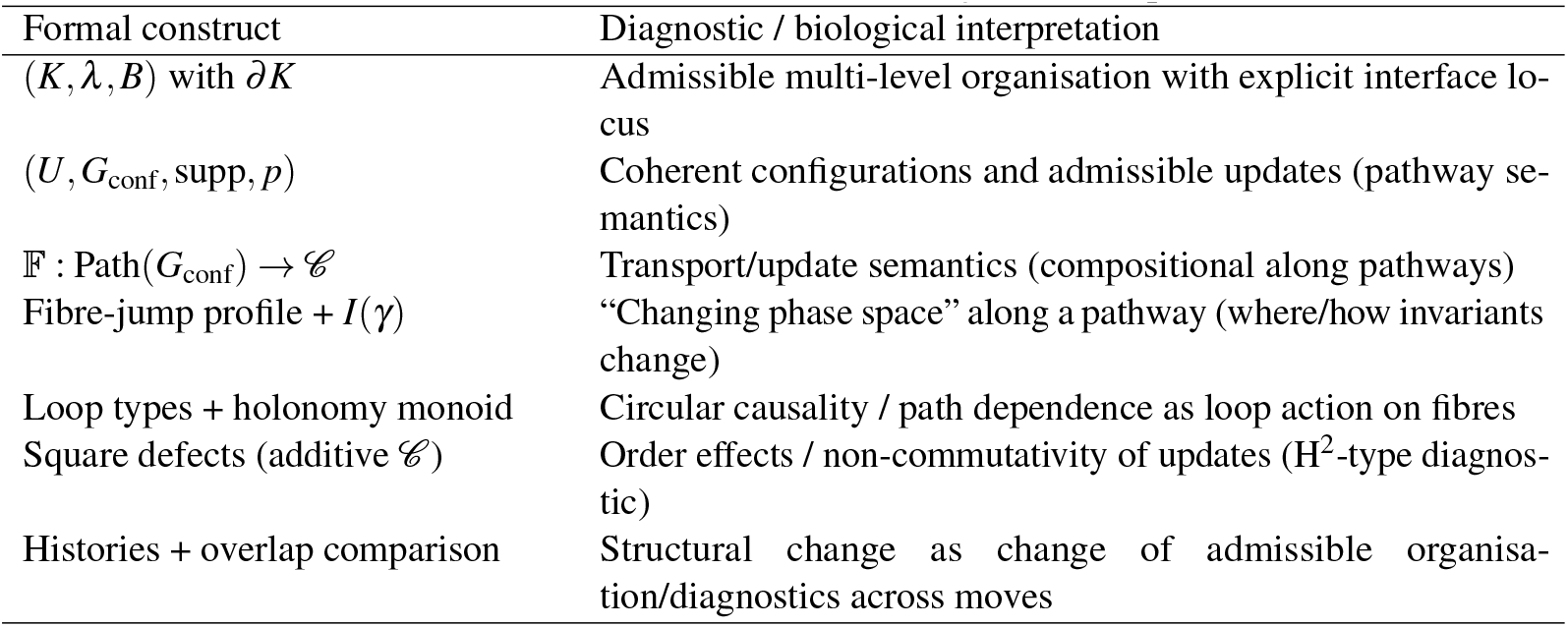
Core formal constructs and their diagnostic interpretation.

**Table 2:**
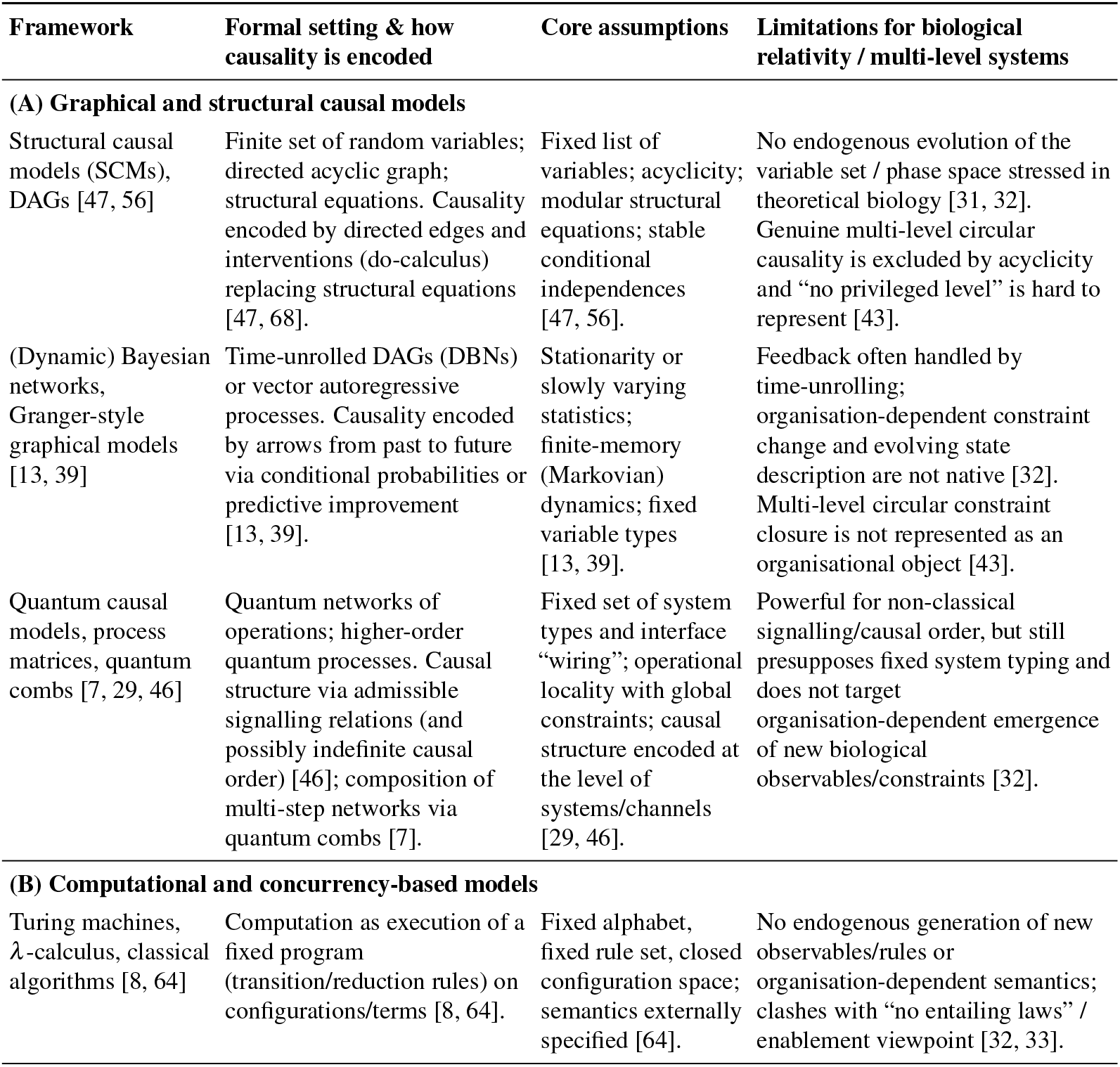
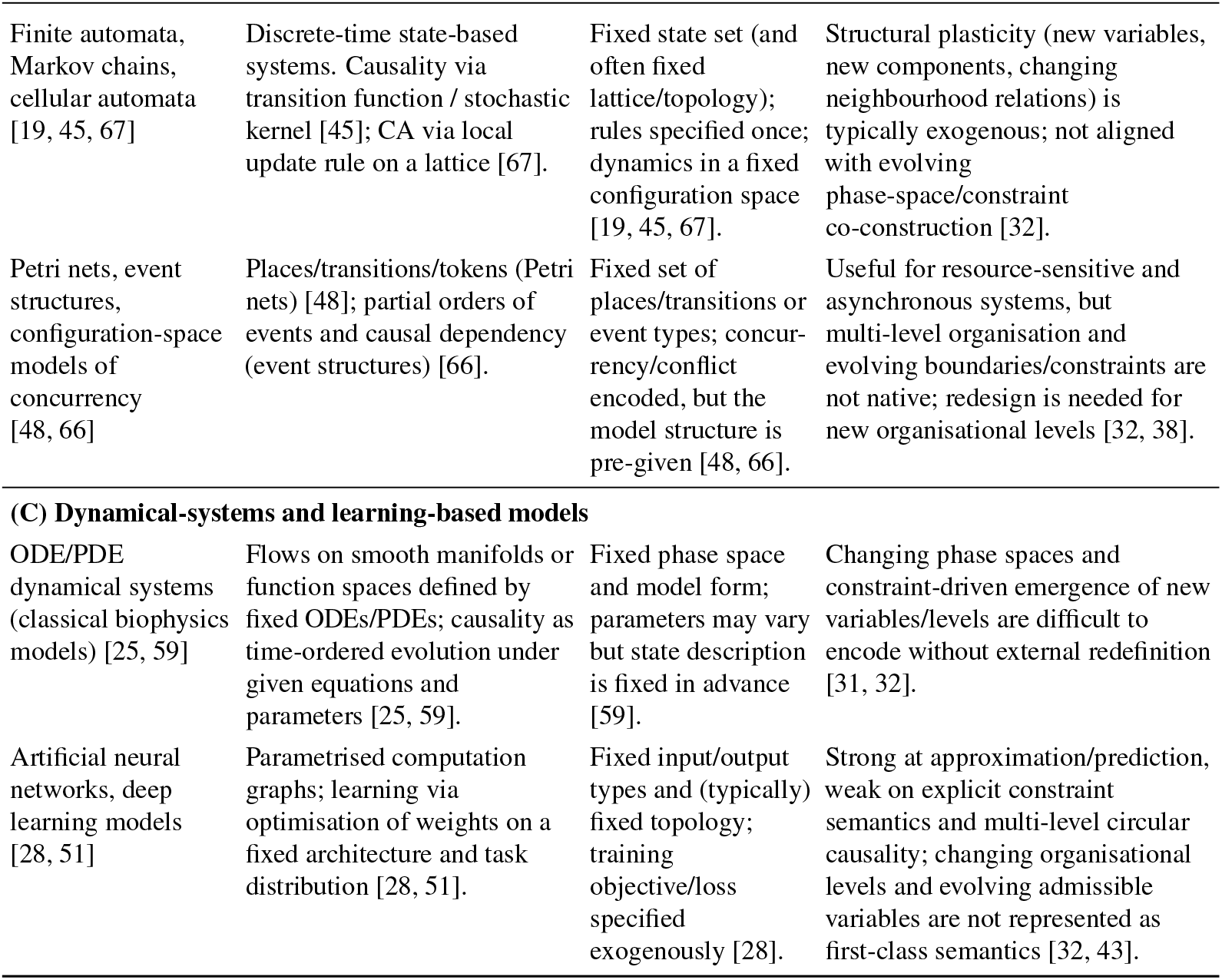
Standard causal and computational frameworks and their limitations for multi-level, constraint-based biological organisation (in the sense of Noble and Longo–Montévil).

**Table 3:**
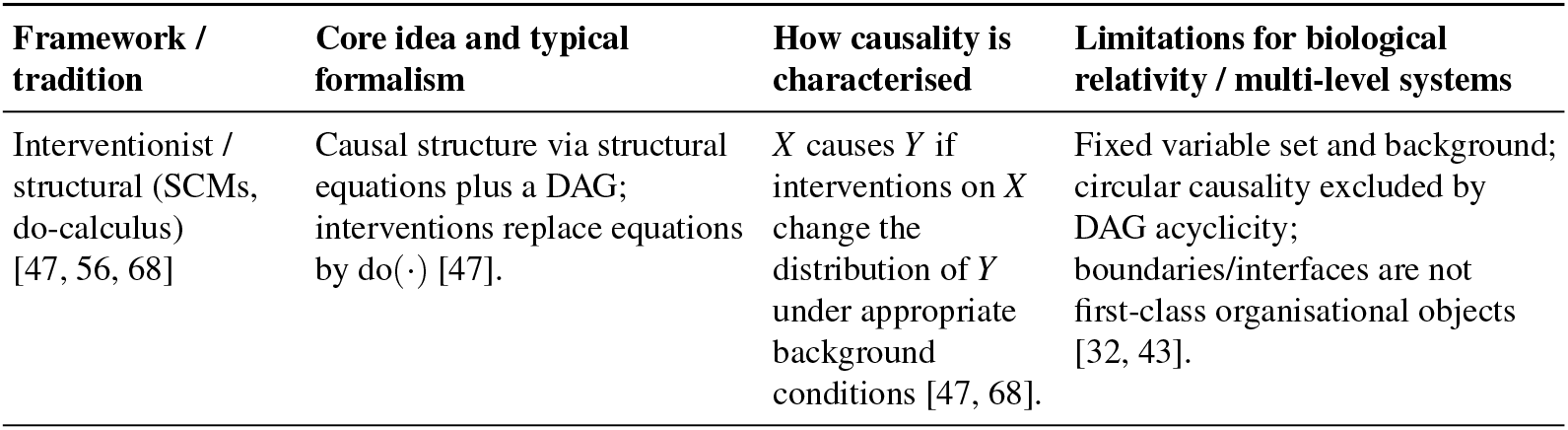
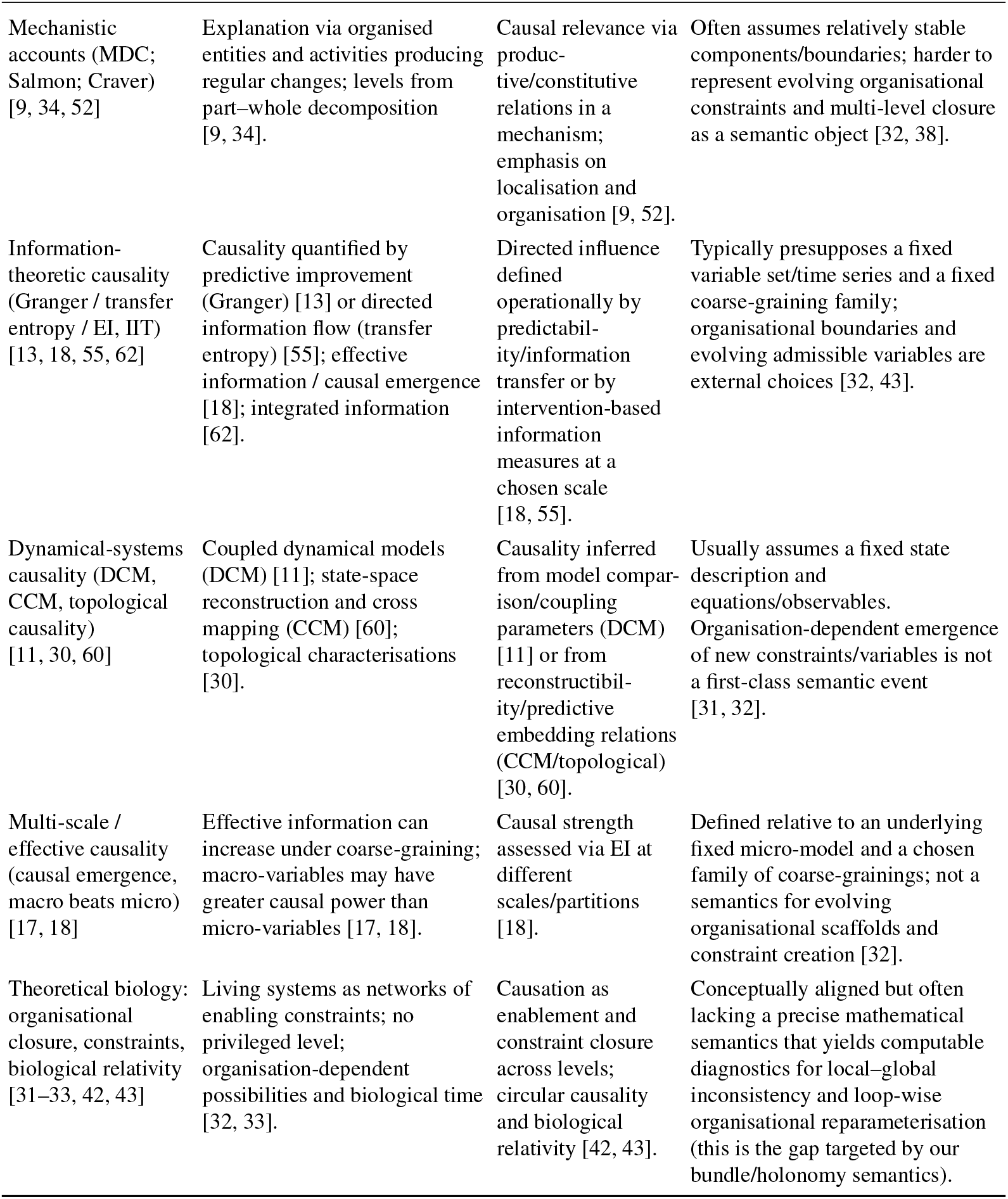
Causal notions and frameworks across disciplines and some limitations for multi-level, constraint-based biological organisation (in the sense of Noble and Longo–Montévil).

| **Framework / tradition** | **Core idea and typical formalism** | **How causality is characterised** | **Limitations for biological relativity / multi-level systems** |
| --- | --- | --- | --- |
| Interventionist / structural (SCMs, do-calculus) [47, 56, 68] | Causal structure via structural equations plus a DAG; interventions replace equations by $\text{do}(\cdot)$ [47]. | $X$ causes $Y$ if interventions on $X$ change the distribution of $Y$ under appropriate background conditions [47, 68]. | Fixed variable set and background; circular causality excluded by DAG acyclicity; boundaries/interfaces are not first-class organisational objects [32, 43]. |

**Table 3 (continued)**
| **Framework / tradition** | **Core idea and typical formalism** | **How causality is characterised** | **Limitations for biological relativity / multi-level systems** |
| --- | --- | --- | --- |
| Mechanistic accounts (MDC; Salmon; Craver) [9, 34, 52] | Explanation via organised entities and activities producing regular changes; levels from part-whole decomposition [9, 34]. | Causal relevance via productive/constitutive relations in a mechanism; emphasis on localisation and organisation [9, 52]. | Often assumes relatively stable components/boundaries; harder to represent evolving organisational constraints and multi-level closure as a semantic object [32, 38]. |
| Information-theoretic causality (Granger / transfer entropy / EI, IIT) [13, 18, 55, 62] | Causality quantified by predictive improvement (Granger) [13] or directed information flow (transfer entropy) [55]; effective information / causal emergence [18]; integrated information [62]. | Directed influence defined operationally by predictability/information transfer or by intervention-based information measures at a chosen scale [18, 55]. | Typically presupposes a fixed variable set/time series and a fixed coarse-graining family; organisational boundaries and evolving admissible variables are external choices [32, 43]. |
| Dynamical-systems causality (DCM, CCM, topological causality) [11, 30, 60] | Coupled dynamical models (DCM) [11]; state-space reconstruction and cross mapping (CCM) [60]; topological characterisations [30]. | Causality inferred from model comparison/coupling parameters (DCM) [11] or from reconstructibility/predictive embedding relations (CCM/topological) [30, 60]. | Usually assumes a fixed state description and equations/observables. Organisation-dependent emergence of new constraints/variables is not a first-class semantic event [31, 32]. |
| Multi-scale / effective causality (causal emergence, macro beats micro) [17, 18] | Effective information can increase under coarse-graining; macro-variables may have greater causal power than micro-variables [17, 18]. | Causal strength assessed via EI at different scales/partitions [18]. | Defined relative to an underlying fixed micro-model and a chosen family of coarse-grainings; not a semantics for evolving organisational scaffolds and constraint creation [32]. |
| Theoretical biology: organisational closure, constraints, biological relativity [31–33, 42, 43] | Living systems as networks of enabling constraints; no privileged level; organisation-dependent possibilities and biological time [32, 33]. | Causation as enablement and constraint closure across levels; circular causality and biological relativity [42, 43]. | Conceptually aligned but often lacking a precise mathematical semantics that yields computable diagnostics for local–global inconsistency and loop-wise organisational reparameterisation (this is the gap targeted by our bundle/holonomy semantics). |

This paper proposes a first step: a *constraint semantics* for multi-level biological dynamics that is faithful to the theoretical-biology viewpoint while remaining operational and mathematically sound [23, 32, 36–38]. The semantic move is to treat *organisation* as a stratified combinatorial bundle (a stratified simplicial complex equipped with an explicit interface/boundary subcomplex), and to treat *functionality* (computation) as transport/rewriting in attached local state spaces (fibres). In this semantics, structure and computation are not separable layers: the organisational scaffold constrains which transports are admissible, while transport along fibres realises organisational pathways. The framework is designed to support theoretical-biology themes—including enablement, path dependence, and internally generated notions of biological time—without claiming to resolve the full programme of changing phase spaces in one stroke [31–33, 35]. Our aim here is deliberately modest: we do not claim to settle these foundational questions in theoretical biology, but rather to provide a *precise mathematical semantics* in which such notions acquire operational meaning. In this sense, our proposal is aligned with existing attempts to give a rigorous account of biological organisation rather than just of its dynamics.

In a complementary mathematical direction, category-theoretic multi-level models (e.g., Memory Evolutive Systems) treat the system’s “state” as a structured categorical object and transitions as functorial transformations, explicitly accommodating heterogeneous components and emergent higher-level objects constructed by limits. Parallel lines of work support the same underlying agenda: a biological system is not adequately described as dynamics on a fixed, homogeneous phase space rather satisfying closure of constraints [10, 35], minimal and autonomous living systems emphasises that cells must be understood as self-maintaining organisations under spatial and energetic constraints[24, 49, 50], geometry-led differential heterogenesis to model the emergence of cortical feature maps and semiotic functions [53, 54], indicates that multilevel pattern-formation and function should be treated in a unified geometric language. Our semantics is designed to sit at this intersection: a step to give a preliminary mathematical structures associated with foundational concepts of theoretical biology.

Organisational dynamics in biology is often described as *history- and organisation-dependent*: the relevant “space of possibilities” is not fixed *a priori*, but is progressively constituted and reshaped by the system’s own ongoing activity—a cluster of ideas articulated as *enablement, changing phase space, extended criticality*, and *closure of constraints*. Concretely, we represent a multilevel organisation by (i) a combinatorial scaffold encoding admissible multilevel configurations and transitions, together with (ii) transport/compatibility data describing how functional states are related across that scaffold; within this semantics, “change of phase space” corresponds to changes in admissible configurations and/or in the associated state-descriptions along a history. In the same language, *circular causality* is encoded as a loop-wise cumulative transport effect (a holonomy/defect), yielding mathematical diagnostics that distinguish coherent regimes with negligible loop effect, persistently twisted regimes with nontrivial loop effect, and unique (rare) *structural reconfiguration* events in which the organisational scaffold itself changes (as opposed to mere reparameterisation on a fixed scaffold).

The semantics also makes precise a central local–global tension: local compatibility constraints may fail to glue into a globally consistent state, and the resulting obstruction/holonomy signatures yield diagnostics of multi-level inconsistency. This diagnostic stance is conceptually parallel to “local-to-global” obstruction perspectives developed in sheaves, obstruction classes (Čech cohomology in the abelian case, and torsors/descent data in the non-abelian case), and stratified settings[12, 15, 40, 58] Our framework repurposes this local-to-global obstruction/holonomy spine as an organisational diagnostic: interfaces and loops are loci where local multi-level compatibilities may fail to globalise, and the resulting holonomy/defect signatures become mathematical indicators of multi-level inconsistency and path dependence.

We illustrate the semantics on a canonical excitable-cell exemplar: a Hodgkin–Huxley-type spike viewed as a cross-level organisational loop spanning cellular, membrane, and molecular constraints [16, 21, 42]. The aim is deliberately modest: we do *not* re-derive Hodgkin–Huxley equations and do *not* claim to resolve the full theoretical-biology programme on changing phase spaces. Rather, we show how to (i) construct a configuration quiver for an excitable-cell organisational scaffold, (ii) compute loop-specific fibre-jump profiles, and (iii) extract holonomy-based diagnostics that compactly quantify circular causal closure and history dependence. Structural reconfiguration is introduced as a principled semantic category primarily relevant over longer histories (development, adaptation, learning), rather than asserted for the single-spike exemplar [32, 43].

A key conceptual point, emphasised in organisational physiology, is that the “Hodgkin cycle” should not be read as a cycle in a fixed phase space. Our formalism makes this precise: the spike is a loop in the *configuration quiver* (returning to an organisational context), while the lifted evolution in the total space traverses non-isomorphic fibres at fibre-jump edges, so there is no single homogeneous state space in which the entire spike is a closed orbit. Accordingly, circular causality is diagnosed by the induced holonomy endomorphism on the resting fibre—a loop-wise return map capturing history dependence—rather than by periodicity in a pre-given phase portrait [43, 44].

### Contributions

The paper makes the following contributions:

(C1) *Constraint semantics for multilevel organisation:* a stratified organisational scaffold with an explicit interface and heterogeneous fibres, formalising structure–computation inseparability as (scaffold, transport) [32, 35, 38].

(C2) *Configuration-quiver semantics:* coherent configurations and admissible moves, with *fibre jumps* encoding organisation-dependent changes of local state descriptions along pathways [32].

(C3) *Loop diagnostics for circular causality:* holonomy endomorphisms induced by configuration loops, and a regime classification (weak/persistent/structural) connecting loops to persistent reparameterisation and unique (rare) reconfiguration [32, 43].

(C4) *Excitable-cell case study:* a Hodgkin spike as a cross-level organisational loop yielding fibre-jump profiles and holonomy diagnostics without re-deriving mechanistic kinetics [16, 21, 42].

(C5) *Comparative positioning:* crosswalks clarifying how the semantics complements standard causal, computational, and dynamical formalisms by treating interfaces and local–global consistency diagnostically (Section 2) [18, 47, 55, 68].

(C6) *Local–global inconsistency as an obstruction diagnostic*: a classical obstruction/holonomy framing of multi-level incompatibility, in which failures of globalisation across interfaces are detected by loop-wise holonomy and typed order-defect operators.

(C7) *Monolithic co-evolution of organisation and dynamics*: a coupled geometry–transport semantic object (organisational scaffold + fibre transport + loop operators) that formalises structure–function inseparability and makes “changing phase space” operational via fibre-jump profiles and history-dependent reconfiguration

### Outline

Section 2 positions the approach within theoretical and computational biology and summarises why active interfaces, circular causality, and organisation-dependent state descriptions motivate additional semantic structure (including the comparative tables). Section 3 introduces the organisational scaffold, configuration quiver, fibre jumps, and holonomy-based diagnostics. Section 4 develops the excitable-cell exemplar and computes loop-level diagnostics for a spike pathway. Section 5 discusses scope, limitations, and extensions to longer-horizon adaptive reconfigurations and interacting organisational loops.

## 2 Background theory and motivation

### Why a constraint semantics for multi-level biology?

Biological systems are organised across multiple, interacting levels (molecular, membrane, cellular, tissue), where constraints propagate both upward and downward. A central theme in theoretical biology is that such systems are not well represented as separable components evolving in a fixed, pre-given state space. Instead, biological organisation is treated as a structured whole in which (i) structure and function are inseparable, (ii) causation often takes the form of *enablement* (opening or reshaping the space of possibilities rather than entailing a unique trajectory), and (iii) the relevant observables and admissible behaviours may change with the organisation itself [31–33].

In physiology, Noble’s principle of *biological relativity* adds a complementary requirement: there is no privileged causal level. Constraints can propagate downward and upward across levels, closing into *circular causality* (e.g. ion-channel kinetics shape the membrane potential, which shapes channel state transitions, while cellular/tissue constraints modulate both) [42, 43]. Organisational perspectives in philosophy of biology formalise related intuitions through *closure of constraints* and *organisational function*, where higher-level structures constrain lower-level processes while being maintained by them [35, 37, 38].

For this paper, these strands motivate a concrete modelling target: we seek a semantics that (a) treats multi-level organisation as a first-class object, (b) makes circular causal pathways operationally visible, (c) supports the possibility that the effective state description changes along organisational histories, and (d) yields computable diagnostics of when local compatibility fails to assemble into a globally coherent description. We use these desiderata as *design constraints* for the formalism rather than as claims that we fully solve the broader theoretical-biology programme [32, 43].

#### Operational desiderata

We will build a semantics that supports:

- *No privileged level of causation:* causal efficacy distributed across levels (biological relativity) [43].
- *Active interfaces/boundaries:* interfaces are sites where constraints are mediated and can change organisational coupling [37, 42].
- *Enablement and organisation-dependent possibilities:* the admissible behaviours may depend on organisational context, not only on local mechanisms [31, 33].
- *Organisation-dependent state description:* modelling should admit that the effective description of “state” can change along histories [32].
- *Local–global tension:* locally compatible pieces may fail to assemble into a globally consistent description, requiring diagnostic invariants/obstructions [1, 27].
- *Loop-wise diagnostics of circular causality:* circular pathways should induce mathematical signatures (holonomy/obstruction-like) rather than remaining implicit [2, 41, 43].

### Standard Modelling Frameworks

Standard frameworks in causality, computation, and dynamics are powerful and indispensable; our claim is not that they are “wrong”, but that many of them share assumptions that make multi-level organisational closure and active interfaces difficult to represent as first-class semantic objects. A recurring pattern is the presupposition of fixed variable types and a fixed configuration space, together with cycles handled indirectly (for example by time-unrolling or by embedding feedback into a larger Markov state) [11, 13, 25, 47, 68]. When the modelling aim is to represent organisation-dependent constraint change and circular causality across levels, these assumptions become limiting [32, 43].

To keep the discussion systematic, we summarise representative formalisms and their typical assumptions in Tables 2 and 3. These tables should be read as a *crosswalk*: they identify what each tradition captures well, and which of our desiderata it does not natively treat as semantic structure (active interfaces, organisation-dependent state description, and explicit loop diagnostics).

### Local-to-global semantics

A key conceptual template for our approach is *local-to-global* reasoning. In many areas of mathematics and physics, global objects are specified by local data plus compatibility constraints; failures of global coherence are detected by invariants and obstruction classes. For example, principal bundles are assembled from local trivial data glued by transition functions satisfying cocycle conditions; the existence of global sections can fail due to topological obstructions [2, 41]. Similarly, in sheaf/bundle semantics for contextuality, locally consistent measurement assignments may fail to glue to a global assignment, and this failure is diagnosed by obstruction-type invariants [1, 27].

Our use of this template is *biology-first*. We do not import quantum-foundational interpretation. Instead, we treat multi-level organisation as a scaffold of local descriptions with explicit compatibility constraints, and we treat circular organisational pathways as loops that induce cumulative transport on local state spaces. These cumulative effects are captured by holonomy-like endomorphisms, enabling loop-wise diagnostics aligned with biological relativity and circular causality [42, 43]. This sets up the core move of the paper: define a configuration space of coherent multi-level states and analyse loops in this space via transport/holonomy signatures.

### Mathematical primer on Fiber Bundles

We fix a (Hausdorff, paracompact) base space *X* and a typical fibre *F*. A *fibre bundle* over *X* with fibre *F* is a surjection

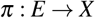

for which there exists an open cover 𝒰 = {*U*_*i*_}_*i*∈*I*_ of *X* and homeomorphisms (local trivialisations)

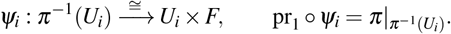

If the allowed changes of fibre coordinates are restricted to a structure group *G* ≤ Homeo(*F*) (or *G* ≤ Diff(*F*) in the smooth case), we speak of a bundle with structure group *G* [20, 57].

#### Transition functions and the Čech 1-cocycle condition

On overlaps *U*_*ij*_ := *U*_*i*_ ∩ *U*_*j*_, the trivialisations differ by a *transition function*

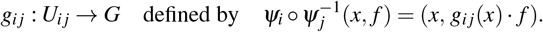

They satisfy the standard identities

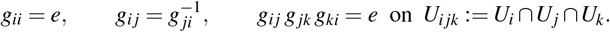

Thus {*g*_*ij*_} is a (generally non-abelian) Čech 1-cocycle with values in *G*. Two cocycles related by a change of trivialisation *h*_*i*_ : *U*_*i*_ → *G*,

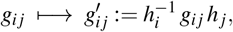

define isomorphic bundles; the corresponding equivalence class is the Čech cohomology class 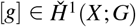 [4, 20].

#### Sections and global coherence

A (local) *section* on *U* ⊆ *X* is a map *s* : *U* → *E* with *π* °*s* = id_*U*_ . Writing *s*_*i*_ for sections on *U*_*i*_, the compatibility (gluing) condition on overlaps is

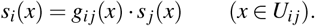

A *global* section is a section on all of *X* . In our semantics, a “globally coherent state” corresponds precisely to a compatible family of local states satisfying these gluing constraints; non-existence of such a global object is interpreted as a *diagnostic* of local–global inconsistency. For principal *G*-bundles, existence of a global section is equivalent to triviality (hence to [*g*] = 0); for general associated bundles one may have sections without full triviality [20, 57].

#### Connection as transport and holonomy as a loop observable

To compare fibre states along paths in *X*, one equips the bundle with a *connection*, whose operational content is *parallel transport*: for a path *γ* : [0, 1] → *X*,

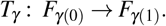

For a loop *γ* based at *x* ∈ *X*, the induced endomorphism

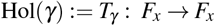

is the *holonomy* of the connection around *γ*; under gauge change it is defined up to conjugation. In smooth settings, this transport is encoded by a principal connection 1-form and curvature (standard references include [26, 41]). In discrete settings (as in many combinatorial models), one can work directly with transport elements on edges and define loop holonomy as an ordered product, mirroring lattice gauge constructions [2, 41].

#### Interface constraints and relative viewpoint (glimpse)

If a distinguished interface/boundary *B* ⊆ *X* is equipped with fixed boundary data (e.g. fixed trivialisation or restricted transitions), then the relevant gluing/obstruction data is naturally formulated *relative* to *B*. At the level of Čech data, this corresponds to cocycles constrained on the subcover induced by *B*, and the resulting obstruction class lives in a relative group (schematically) of the form 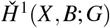 [14]. We use this as a conceptual guide for “interface vs ambient” diagnostics, while keeping the full formal development in Section 3.

From a more structural point of view, our use of organisational bases, fibres, and gluing data can be seen as fitting within a Grothendieck-style semantic picture. Informally, the family of organisational contexts (simplices, strata, interface patches) together with their incidence relations forms a natural “site” of contexts, and the organisational bundles we consider amount to sheaf- or bundle-like assignments of local state spaces over this site. This resonates with Caramello’s topos-theoretic programme of using “toposes as bridges” between different mathematical theories [5, 6]. In our setting, evolving organisational simplicial bundles provide one candidate semantic environment on the geometric side of such a bridge, into which more traditional dynamical, network, or process-theoretic descriptions of the same biological system could in principle be embedded.

#### Biological Mapping

Mathematically, (*X, E, π, F, G*) is standard bundle data; biologically, the intended reading is:

- *X* : an *organisational domain* indexing where/which local descriptions apply (context, locus, organisational condition).
- *F*: a *local state space* (variables/parameters admissible at a locus).
- *g*_*ij*_: *compatibility/constraint maps* translating local descriptions between overlapping contexts.
- global section *s*: a *globally coherent state* compatible with all constraints.
- holonomy Hol(*γ*): the *net effect of traversing a circular pathway*, serving as a loop-wise diagnostic of circular causality and history dependence.

## 3 Evolving Organisational Simplicial Bundles

We formalise a combinatorial–geometric framework for multi-level biological organisation in which (i) the admissible organisational scaffold is represented by a *stratified simplicial complex with explicit interfaces*, and (ii) multi-level pathways (corresponding to biological pathways like glycolysis, Hodgkin cycle, etc) are analysed on a *derived discrete base* (a configuration quiver) on which *transport, fibre jumps, and loop holonomy* are computable. The key distinction between organisational *possibilities/constraints* (the scaffold) and *coherent traversals* of those possibilities (the pathways) provides a mathematical semantics that aligns with the theoretical-biology view that, in evolution and development, the space of admissible configurations is historically produced and reconfigured, rather than fixed in advance by an entailing law [31, 33]. We do not claim to provide a comprehensive mathematical semantics to this programme rather a first step to map their core ideas in a Grothendieck-style formalism.

### Stratified Organisational Scaffold

#### Definition 3.1.

(Organisational strata and pure scaffold). Let *L* be a finite partially ordered set (poset) of organisational levels (e.g. molecular *<* membrane *<* cellular *<* tissue). An *organisational family of strata* is a collection of finite abstract simplicial complexes { *K*_*ℓ*_ }_*ℓ*∈*L*_ such that their vertex sets are pairwise disjoint:

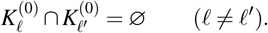

The associated *pure organisational scaffold* is the simplicial complex

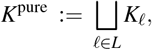

whose vertex set is 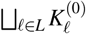 and whose simplices are precisely the simplices of the individual strata *K*_*ℓ*_.

Throughout, vertices and simplices should be read as *organisational descriptors*: they encode patterns, relations, or coarse-grained configurations at different levels, not literal physical points in Euclidean space.

#### Definition 3.2.

(Interface subcomplex and global organisational scaffold). Let { *K*_*ℓ*_ }_*ℓ*∈*L*_ be an organisational family of strata and let *K*^pure^ = ∐_*ℓ*∈*L*_ *K*_*ℓ*_. An *interface subcomplex* is a finite abstract simplicial complex *B* such that:

i. 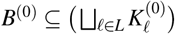
ii. every simplex *τ* ∈ *B* is *mixed-level*: its vertex set meets at least two distinct levels in *L*.

Moreover, we impose the compatibility condition:

(⋆) For every *τ* ∈ *B* and every *ℓ* ∈ *L*, the (possibly empty) vertex subset 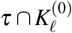 spans a simplex of *K*_*ℓ*_.

The *global organisational scaffold* is the simplicial complex

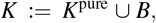

formed by adjoining the interface simplices to the pure scaffold. We refer to *K* as a *stratified organisational simplicial complex*.

#### Definition 3.3.

(Level map, pure/mixed simplices, and organisational boundary). Let *K* = (∐*ℓ*∈*L K*_*ℓ*_) ∪ *B* be a stratified organisational simplicial complex.

a. (*Level map*.) For each vertex *v* ∈ *K*^(0)^ there is a unique *ℓ* ∈ *L* such that 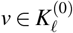. Define *λ* : *K*^(0)^ → *L* by *λ* (*v*) = *ℓ*. For a simplex *σ* ∈ *K* with vertex set {*v*_0_, …, *v*_*k*_}, set *λ* (*σ* ) := {*λ* (*v*_0_), …, *λ* (*v*_*k*_)} ⊆ *L*. We call *σ pure* if |*λ* (*σ* )| = 1 and *mixed* otherwise.
b. (*Interface boundary*.) Fix two levels *ℓ < ℓ*^*′*^ in *L*. The *interface boundary of K*_*ℓ*_ *towards ℓ*^*′*^ is the subcomplex

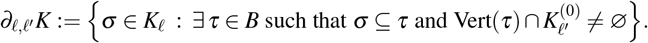

The *organisational boundary* of *K* is the union *∂K* := ∪ _*ℓ<ℓ*_*′ ∂*_*ℓ,ℓ*_*′ K*.

The boundary *∂K* is the combinatorial locus where cross-level constraints are expressed. In the sequel, transport effects associated with between-level coupling and multi-level circular dependence are naturally localised near *∂K*.

#### Remark 1

(Bidirectionality is a semantic requirement). *In stratified-topology settings, one often uses exit/entrance path categories which privilege a direction of the filtration. Here organisational levels are not ordered by an underlying Euclidean dimension, and reciprocal inter-level influence (e*.*g. cell* ↔ *membrane* ↔ *molecular) is central [31, 43]. Accordingly, we do* not *impose exit/entrance constraints on paths. Instead, bidirectionality is enforced in the configuration quiver construction below (Definition 3.9)*.

### Reconfiguration of the scaffold and irreversible time

Organisational change is represented by specifying a class of admissible *reconfiguration moves* between scaffolds and composing these moves into realised histories, rather than postulating an entailing evolution law of the form *t* → (*K*_*t*_, *λ*_*t*_, *B*_*t*_).

#### Definition 3.4.

(Move system of organisational scaffolds). Fix a finite level poset *L*. Let Org(*L*) denote the class of stratified organisational simplicial complexes (*K, λ, B*) satisfying Definitions 3.1–3.3. An *organisational reconfiguration move* is a map

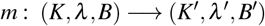

given by a finite modification of the simplicial data such that (*K*^*′*^, *λ*^*′*^, *B*^*′*^) ∈ Org(*L*). The admissible moves define a directed multigraph on Org(*L*); a directed path in this multigraph is an *organisational history*.

#### Definition 3.5.

(Elementary organisational reconfiguration moves). An admissible move *m* : (*K, λ, B*) → (*K*^*′*^, *λ*^*′*^, *B*^*′*^) is *elementary* if it is of one of the following types:

(E1) *Within-stratum update*. For some *ℓ* ∈ *L*, replace *K*_*ℓ*_ by 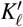 on the same vertex set 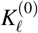 by adding/removing finitely many simplices, leaving all other strata unchanged and keeping *B* fixed.

(E2) *Interface update*. Replace *B* by *B*^*′*^ obtained by adding/removing finitely many mixed simplices, keeping all strata fixed, and preserving condition (⋆) in Definition 3.2.

General reconfigurations are finite compositions of elementary moves.

#### Definition 3.6.

(Organisational histories and irreversible order). A *(finite) organisational history* is a composable sequence of admissible moves

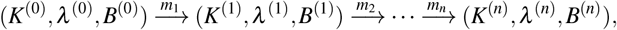

with each (*K*^(*i*)^, *λ* ^(*i*)^, *B*^(*i*)^) ∈ Org(*L*). An *(infinite) organisational history* is a countable composable sequence indexed by ℕ. The index order induces an irreversible order: once a move has occurred, it is part of the realised history.

#### Remark 2

(Internal irreversible axis). *Organisational histories provide the first component of internal biological time: an intrinsic, discrete irreversible axis recording how the admissible scaffold itself changes, without presupposing a pre-given trajectory in a fixed phase space [31, 33]*.

### Fibres, heterogeneity, and fibre jumps

The organisational scaffold specifies admissible *patterns*. We attach to each pattern a space of compatible local states or models.

#### Definition 3.7.

(Fibre assignment on simplices). Let Simp(*K*) denote the set of simplices of *K*. An organisational fibre assignment is a map

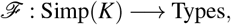

assigning to each simplex *σ* ∈ Simp(*K*) a semantic type ℱ (*σ* ), interpreted as the space of admissible local states or models compatible with *σ*.

#### Remark 3

(Heterogeneity of fibres). *The assignment* ℱ *is generally heterogeneous in two independent senses:*

i. Type heterogeneity: *different simplices may carry different kinds of state/model spaces (finite sets, probability simplices, manifolds, vector spaces, parameter families)*.
ii. Dimensional heterogeneity: *even after choosing a common representation, the effective dimension or complexity of fibres may vary with organisational level and pattern*.

#### Remark 4

(Heterogeneity assumptions in the core formalism). *We model heterogeneity primarily* across organisational layers *(and across interface-mediated supports) by allowing distinct fibre-types across strata and by permitting fibre jumps along admissible configuration updates. For tractability of transport and diagnostics in this paper, we do not attempt full “arbitrary variation everywhere” within a fixed stratum; such within-stratum categorical heterogeneity naturally calls for a site/topology of refinements and fibred-category/stack semantics (with projective limits/descent) to manage multiple interacting populations. We defer that refinement to the Discussion*.

#### Definition 3.8.

(Realised fibre category). Fix a category 𝒞 and a realisation functor *R* : Types → 𝒞 . The realised fibre of a simplex *σ* ∈ Simp(*K*) is the object

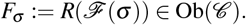

#### Remark 5

(Notation). *We use* ℱ *for the simplex-level fibre assignment* Simp(*K*) → Types *and* 𝔽 *for the discrete organisational bundle functor* Path(*G*_conf_) → 𝒞 . *The realised fibre over a simplex σ is F*_*σ*_ := *R*(ℱ (*σ* )).

#### Remark 6

(Structure category and refined fibre-jumps). *For the conceptual development and diagnostics in this paper it is convenient to work in a fixed concrete category. Unless stated otherwise, we take* 𝒞 = **FinVect**_ℝ_. *If one needs “equidimensional but non-isomorphic” fibres (Type II below), one works instead in a* structured *fibre category (e*.*g. representations, inner-product spaces, graded/decomposed spaces) where additional structure is part of the isomorphism notion*.

### Organisational configurations and configuration quiver

A pathway is a sequence of *organisational configurations* that traverse and coordinate multiple strata and interface simplices.

#### Definition 3.9.

(Organisational configurations and derived base). Let *K* be a stratified organisational simplicial complex. A *derived organisational base* consists of data (*U, G*_conf_, supp, *p*) where:

a. *U* is a set of *organisational configurations*;
b. *G*_conf_ is a directed multigraph (a quiver) with vertex set *U* ;
c. supp : *U* → 𝒫(Simp(*K*)) assigns a finite, nonempty, face-closed set of supporting simplices to each *u* ∈ *U* ;
d. *p* : *U* → Simp(*K*) chooses a representative simplex *p*(*u*) ∈ supp(*u*).

These data satisfy:

(C1) *Subcomplex support*. If *σ* ∈ supp(*u*) and *τ* ⊆ *σ* is a face, then *τ* ∈ supp(*u*).

(C2) *Locality of transitions*. For each edge *e* : *u* → *v* in *G*_conf_, there exists a finite chain of simplices *σ*_0_, …, *σ*_*m*_ ∈ supp(*u*) ∪ supp(*v*) with *σ*_0_ = *p*(*u*), *σ*_*m*_ = *p*(*v*), and each step is a face/coface move (*σ*_*j*_ ⊆ *σ*_*j*+1_ or *σ*_*j*+1_ ⊆ *σ*_*j*_).

(C3) *Interface compatibility*. If an edge *e* : *u* → *v* changes level support (Definition 3.10), then (supp(*u*) ∪ supp(*v*)) ∩*B*≠ ∅.

(C4) *Bidirectional cross-level closure*. If *e* : *u* → *v* is cross-level (i.e. Λ(*u*)≠ Λ(*v*)), then there exists an admissible reverse edge *e*^op^ : *v* → *u*. (No invertibility of fibre maps is assumed.)

#### Definition 3.10.

(Level support). For *u* ∈ *U*, define the *level support*

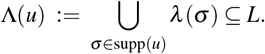

#### Lemma 3.11

(Interface visitation along cross-level paths). *Let u*_0_ → *u*_1_ → · · · → *u*_*k*_ *be a directed path in G*_conf_. *If* Λ(*u*_0_)≠ Λ(*u*_*k*_), *then there exists i* ∈ {0,…, *k*} *such that* supp(*u*_*i*_) ∩*B*≠ ∅.

*Proof*. If Λ(*u*_0_) ≠ Λ(*u*_*k*_), then some edge along the path changes level support. By axiom (C3) of Definition 3.9, any such cross-level edge is interface-mediated, hence its endpoints’ supports intersect *B*.

#### Remark 7

(Canonicity of the configuration quiver). *The derived base* (*U, G*_conf_, supp, *p*) *is* canonical relative to the modelling choices *(what counts as a configuration, which local updates are admitted, and how supports are assigned). We do not claim that G*_conf_ *is uniquely determined by the simplicial scaffold K alone; rather, K constrains admissibility and locality of transitions*.

#### Remark 8

(Relation to stratified geometry and constructible semantics). *Our stratified scaffold/interface construction is a finite combinatorial analogue of classical stratification theory and stratified mappings (Thom–Mather, Verona), and it interfaces naturally with constructible/exit-path viewpoints on stratified phenomena; we cite these works only as mathematical context and do not impose their monotone exit/entrance directionality*.

#### Remark 9

(Relation to Thom stratified morphisms; key differences). *In the Thom–Mather programme one studies a single stratified morphism f* : *X* → *Y (with local triviality over strata) and extracts monodromy/vanishing-cycle holonomies around singular values [22, 61, 63, 65]. Our holonomy diagnostics are analogous in spirit, but the semantic base here is an* operational configuration quiver *encoding admissible multi-level updates with explicit bidirectionality (no privileged causal direction), we allow non-invertible transport (monoid-valued holonomy), and we treat scaffold reconfiguration moves along histories as first-class events, reflecting “historically produced” configuration spaces in theoretical biology*.

#### Why the “Hodgkin cycle” is not a cycle in a fixed phase space

In the classical dynamical-systems sense, the spike is not a closed orbit in a single, pre-stated phase space. In our semantics, the object that is genuinely cyclic is the *organisational* loop *γ*_spike_ in the configuration quiver *G*_conf_: it records a return to a comparable organisational context, not a return to an identical microstate in a fixed coordinate system. Along *γ*_spike_, fibre-jump edges (Definition 3.15) make explicit that adjacent configurations may carry non-isomorphic fibres, so the effective state description changes during the traversal. Consequently, after passing to the total space *E*_spike_ = ∐_*u*_ {*u*} × 𝔽(*u*), the lifted “spike trajectory” is a stratified zigzag across different fibre components and gluing zones, rather than a continuous closed curve in a single state space. The appropriate loop observable is therefore the holonomy endomorphism *H*_spike_ = 𝔽(*γ*_spike_), which returns states to the resting fibre up to a nontrivial return map (refractoriness/memory), rather than enforcing periodicity in a fixed phase portrait.

### Discrete organisational bundles and fibre-jump diagnostics

#### Definition 3.12.

(Path category). The *path category* Path(*G*_conf_) has objects *U* and morphisms finite directed paths in *G*_conf_, with composition by concatenation.

#### Definition 3.13.

(Discrete organisational bundle). A *discrete organisational bundle* over *G*_conf_ is a functor

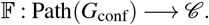

#### Remark 10

(Bundle as functor, no local triviality assumed). *In this paper a “discrete organisational bundle” is* by definition *a functor* 𝔽 : Path(*G*_conf_) → 𝒞 . *We do not assume any topology on G*_conf_ *nor any local triviality in the classical sense; the functorial viewpoint is the appropriate Grothendieck-style semantics for transport on a directed base*.

#### Definition 3.14.

(Support-composite fibres (multi-simplex fibres)). Assume 𝒞 is monoidal, and fix a monoidal product ⊙ (e.g. ⊕ in **FinVect**_ℝ_). A discrete organisational bundle 𝔽 is *support-compatible* with the simplex-level realised fibres *F*_*σ*_ if for each configuration *u* ∈ *U*,

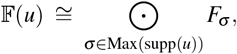

where Max(supp(*u*)) denotes the inclusion-maximal simplices in supp(*u*). This expresses that configurations are multi-level, multi-simplex aggregates rather than single-simplex states.

#### Definition 3.15.

(Fibre jumps and changing phase space). An edge *e* : *u* → *v* in *G*_conf_ is a *fibre jump* if 𝔽(*u*) ≇ 𝔽(*v*) in 𝒞 . A path *γ* has a *fibre-jump profile* given by the positions of its fibre-jump edges.

#### Remark 11

(Dimension diagnostic in **FinVect**_ℝ_). *If* 𝒞 = **FinVect**_ℝ_, *then* 𝔽(*u*) ≅ 𝔽(*v*) *if and only if* dim 𝔽(*u*) = dim 𝔽(*v*). *Hence fibre jumps are detected by the integer-valued dimension profile d*(*u*) = dim 𝔽(*u*).

### Classification of fibre jumps and phase-space changing paths

#### Definition 3.16.

(Types of fibre jumps). Let *e* : *u* → *v* be an edge in *G*_conf_. Write *F*_*u*_ := 𝔽(*u*), *F*_*v*_ := 𝔽(*v*). We say that *e* is a fibre jump of:

i. *Type I (dimensional jump):* dim *F*_*u*_≠ dim *F*_*v*_ (in **FinVect**_ℝ_).
ii. *Type II (structured-type jump at fixed dimension):* dim *F*_*u*_ = dim *F*_*v*_ but *F*_*u*_ ≇ *F*_*v*_ in a *structured* fibre category (Remark 6).
iii. *Type III (irreversible transport at fixed object type): F*_*u*_ ≅ *F*_*v*_ but the transport map 𝔽(*e*) is not an isomorphism.

#### Remark 12

(Fibre invariants and phase-space profiles). *To make “changing phase space” precise beyond dimension alone, fix a family of isomorphism-invariants I* = (*I*_1_, …, *I*_*r*_) *on objects of* 𝒞 *(or on the structured fibre category used in a given application). For a configuration u* ∈ *U, write*

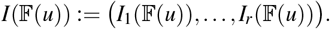

*For a directed path γ* : *u*_0_ → · · · → *u*_*k*_ *in G*_conf_, *define its* phase-space profile *as the sequence*

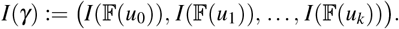

*Every fibre-jump edge e* : *u* → *v forces* 𝔽(*u*) ≇ 𝔽(*v*), *hence I*_*j*_(𝔽(*u*)) ≠ *I*_*j*_(𝔽(*v*)) *for at least one j. Thus fibre-jump profiles localise* where *the invariants change, while I*(*γ*) *records* how *they change*.

#### Definition 3.17

(Phase-space stable vs. phase-space changing paths). A path *γ* is *phase-space stable* if all fibres along *γ* are mutually isomorphic and all edge maps are isomorphisms. Otherwise, *γ* is *phase-space changing*.

#### Remark 13

(Fibre jumps vs. structural reconfiguration). *It is important to distinguish two logically independent kinds of change*.

a. Structural reconfiguration of the scaffold *occurs when a move in the history changes the organisational simplicial complex itself, e*.*g. by adding or removing simplices in K or altering the interface B. This is captured by the move system and organisational histories (Definitions 3.4–3.6)*.
b. Phase-space change along a fixed scaffold *occurs when, for a fixed* (*K, λ, B*) *and derived base* (*U, G*_conf_, *p*), *the bundle* 𝔽 *assigns non-isomorphic fibres to adjacent configurations, i*.*e. along fibre-jump edges in the sense of Definition 3.15*.

*Structural reconfiguration changes which configurations and pathways are even admissible, whereas fibre jumps change the state/model spaces available along a given admissible pathway. In particular, a pathway may exhibit changing phase space (non-trivial fibre-jump profile) without any structural change in the underlying scaffold, and conversely a structural reconfiguration may occur without any fibre jumps along a given local trajectory*.

#### Proposition 3.18

(Characterisation of phase-space stability). *A directed path γ is phase-space stable if and only if*

i. *all fibres along γ are mutually isomorphic as objects of* 𝒞, *and*
ii. *all edge maps along γ are isomorphisms in* 𝒞.

*Equivalently, γ is phase-space changing if and only if it contains at least one fibre jump (Type I or Type II) or at least one non-invertible transport step (Type III), in the sense of your fibre-jump taxonomy*.

*Proof*. Immediate from the definitions: phase-space stability is precisely the conjunction of (isomorphism of objects along the path) and (invertibility of the transport morphisms).

#### Lemma 3.19

(Dimension profile as a minimal fibre-jump diagnostic). *Assume* 𝒞 = **FinVect**_ℝ_ *and let d* : *U* → ℕ *be the dimension profile d*(*u*) := dim_ℝ_ 𝔽(*u*). *Then:*

a. *every Type I fibre-jump edge e* : *u* → *v satisfies d*(*u*)≠ *d*(*v*), *hence is detected by d;*
b. *fibre jumps at constant dimension (e*.*g. Type II in a structured fibre category) cannot be detected by d alone and require refined invariants (cf. Remark 12)*.

*Proof*. In **FinVect**_ℝ_, isomorphism of objects is equivalent to equality of dimension; the claims follow.

#### Remark 14

(Biological and physical meaning of fibre jumps). *Fibre jumps formalise a local change of phase space along a pathway: the space of admissible states or models changes because the organisational context has changed, typically by moving between levels or across the interface. In the biological interpretation, a fibre jump may correspond to:*

- *the recruitment of new degrees of freedom (e*.*g. opening of an ion channel that brings a new mode of ionic flow into play);*
- *coarse-graining or loss of degrees of freedom (e*.*g. passing from a detailed molecular configuration to an effective membrane conductance);*
- *a qualitative change in which variables are relevant (e*.*g. switching from a description in terms of concentrations to one in terms of effective rates or thresholds)*.

*In contrast, edges that are not fibre jumps encode transport within a fixed local phase space. Fibre-jump profiles therefore provide a concrete diagnostic of how changing organisational context induces changing phase spaces along biological trajectories, in the sense advocated by Longo and Montévil [31], Longo et al. [33]*.

#### Remark 15

(Biological reading of phase-space changing paths). *In the biological interpretation, Type I and Type II jumps mark points where the number or nature of effective degrees of freedom changes, for example when an additional slow variable becomes relevant or when a detailed molecular description is replaced by an effective conductance model. Type III jumps capture dissipative, irreversible transformations inside a fixed phase space (for example, projections that lose information). Phase-space changing paths therefore provide a direct diagnostic of the “changing phase space” behaviour emphasised in theoretical biology: as the organisational context changes along the pathway, the admissible local models and degrees of freedom change with it, rather than being fixed once and for all*.

#### Remark 16

(Changing phase space vs scaffold reconfiguration). *In our framework there are two distinct ways in which “phase space” can change:*

i. Scaffold reconfiguration, *encoded at the level of organisational histories* (*K*^(*i*)^, *λ* ^(*i*)^, *B*^(*i*)^), *alters the space of admissible organisational patterns and hence the potential repertoire of future configurations and bundles*.

i. Fibre jumps along a fixed scaffold, *encoded by non-isomorphic fibres* 𝔽(*u*) ≇ 𝔽(*v*) *along an edge u* → *v in a given configuration graph G*_conf_, *alter the local space of admissible states or models within a fixed organisational context*.

*Both phenomena correspond to “changing phase space” in the sense of theoretical biology [33], but they operate at different structural levels: (i) changes the geometry of the organisational scaffold, whereas (ii) changes the fibre-level description over a fixed portion of that scaffold*.

### Loop structure: intra-level and cross-level cycles

#### Definition 3.20.

(Loop types in the configuration graph). A *loop* based at *u*_0_ ∈ *U* is a finite directed cycle *u*_0_ → *u*_1_ → · · · → *u*_*k*_ → *u*_0_ in *G*_conf_. Its *level support* is 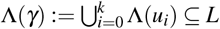. The loop is *stratum-pure* if Λ(*γ*) = { *ℓ* } and *cross-level* if | Λ(*γ*) | ≥ 2. A cross-level loop is an *interface loop* if supp(*u*_*i*_) ∩ *B* ≠ ∅ for some *i*.

#### Remark 17

(Biological interpretation). *Stratum-pure loops correspond to recurrent patterns confined to a single level (e*.*g. a local channel-gating cycle). Cross-level loops formalise multi-level circular causality (levels reciprocally constrain/enable one another), a central theme of biological relativity [43]. Interface loops isolate the boundary-mediated component of cross-level feedback*.

#### Definition 3.21

(Multi-level circular causality loop with phase-space change). A loop *γ* is a *multi-level circular causality loop with phase-space change* if it is cross-level, visits the interface, has nontrivial holonomy (Definition 3.22), and contains at least one fibre jump (Definition 3.16).

#### Remark 18

(Hierarchy of loop types). *Within this framework we obtain a natural hierarchy:*

a. *Stratum-pure loops without fibre jumps and with trivial holonomy encode recurrent patterns inside a single level and a fixed phase space*.
b. *Cross-level loops without fibre jumps but with nontrivial holonomy encode multi-level cycles in which the state is transformed in a path-dependent way, but the effective phase space remains fixed*.
c. *Multi-level circular causality loops with phase-space change (Definition 3.21) encode cycles in which different levels constrain and enable one another in a recurrent pattern, and the very space of admissible local models changes along the loop*.

*In the biological reading, the last class provides a mathematically precise counterpart to the notion of circular causality across organisational boundaries, in which cycles do not simply “run around a fixed phase portrait” but reorganise which variables and degrees of freedom are available*.

### Holonomy, path dependence, and diagnostic regimes

#### Definition 3.22.

(Holonomy monoid at a configuration). Let 𝔽 : Path(*G*_conf_) → 𝒞 be a discrete organisational bundle and fix *u* ∈ *U* . The *holonomy monoid* at *u* is the submonoid of End_𝒞_ (𝔽(*u*)) generated by 𝔽(*γ*) over all loops *γ* based at *u*. If all edge maps along a region are isomorphisms, holonomy takes values in Aut_𝒞_ (𝔽(*u*)).

#### Definition 3.23.

(Holonomy representation and *H*^1^-type diagnostic). Fix a connected subgraph *H* ⊆ *G*_conf_ and basepoint *u*_0_. If all edge maps along *H* are isomorphisms, then 𝔽 induces a group homomorphism

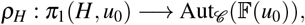

well-defined up to conjugation. Here *π*_1_(*H, u*_0_) denotes the fundamental group of the geometric realisation of the *underlying undirected* 1-complex of *H*. The conjugacy class of *ρ*_*H*_ is the *H*^1^-type holonomy diagnostic on *H*. Here *π*_1_(*H, u*_0_) denotes the fundamental group of the geometric realisation of the *underlying undirected* 1-complex of *H*.

#### Definition 3.24.

(Order effects and an *H*^2^-type diagnostic on squares). Assume 𝒞 is additive. Let 𝒮 be a chosen set of directed squares (two-step alternative composites)

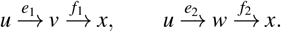

Fix a region on which fibres are pairwise isomorphic and choose trivialisations 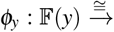 *F*_⋆_ for *y* ∈ {*u, v, w, x*}. Define the *order-defect endomorphism* on the typical fibre by

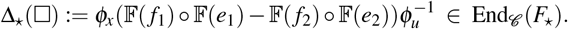

Under a change of trivialisation *φ*_*y*_ ↦ *g*_*y*_ ° *φ*_*y*_ with *g*_*y*_ ∈ Aut_𝒞_ (*F*_⋆_), Δ_⋆_(□) transforms by conjugation; hence its conjugacy class (and any class-function statistic thereof) is gauge-invariant. We treat Δ_⋆_ as an *H*^2^-type obstruction diagnostic; in the present work we state this layer formally but do not attempt full quantitative estimation in examples.

#### Remark 19

(Square-enriched 2-complex). *The pair* (*H*, 𝒮 ) *should be regarded as a square-enriched directed* 2*-complex:* 𝒮 *specifies the 2-cells on which order comparisons are evaluated; without such 2-cells, no H*^2^*-type order diagnostic is available*.

#### Definition 3.25.

(Holonomy regimes). Let *H* ⊆ *G*_conf_.

i. *Weak holonomy:* 𝔽(*γ*) = id for every loop *γ* in *H*.
ii. *Persistent holonomy:* there exist loops with nontrivial 𝔽(*γ*) whose qualitative signatures persist under small admissible perturbations.
iii. *Structural holonomy:* along an organisational history, the holonomy diagnostics change in a way that requires a scaffold reconfiguration (not a small deformation of fibre maps).

### Internal biological time as interaction of histories and loops

The formalism carries two intrinsic temporal structures. First, an organisational history (Definition 3.6) provides an irreversible compositional order of scaffold reconfigurations. This realises an *internal irreversible time axis*: there is a distinguished origin and a partial order of stages, but no external time parameter prescribing an entailing trajectory in a fixed phase space [33].

Second, at each fixed stage of a realised history, the configuration graph *G*_conf_ and its loops support recurrent execution. For a fixed scaffold and configuration graph, the loop structure of *G*_conf_ therefore induces a *circular axis* of time: the action of loops on fibre states via the functor 𝔽 encodes biological rhythms and cycles (for example, repeated spikes, oscillations, or metabolic cycles).

#### Remark 20

(Two-axis internal time). *In combination, the irreversible axis of organisational histories and the circular axis of loop actions yield an internal two-axis structure of biological time. The irreversible axis tracks the production and reconfiguration of the organisational scaffold and of the repertoire of admissible pathways, while the circular axis records the action of recurrent patterns on local state spaces at each stage. This matches the idea, articulated in Longo and Montévil [31] and related work, that biological time is not a single scalar parameter but a richer structure emerging from the organisation itself. In our setting, this structure is encoded by the categorical data of histories, configuration quivers, and their associated bundles, rather than being imposed from outside*.

### Reconfigurable bundles along a history and structural-change criterion

#### Definition 3.26.

(Reconfigurable organisational bundle). Let (*K*^(0)^, *λ* ^(0)^, *B*^(0)^) → · · · → (*K*^(*m*)^, *λ* ^(*m*)^, *B*^(*m*)^) be an organisational history. For each stage *i*, choose a derived base 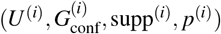 and a bundle 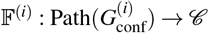. For each adjacent pair *i* → *i* + 1, let *H*^(*i,i*+1)^ be a common overlap subgraph on configurations persisting across the move. A *reconfiguration comparison* is a natural isomorphism

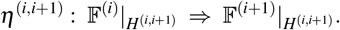

#### Proposition 3.27

(Invariance under reconfiguration comparison on overlaps). *Let* 𝔽^(*i*)^ *and* 𝔽^(*i*+1)^ *be consecutive stage bundles and suppose a reconfiguration comparison (natural isomorphism) is given on the overlap subgraph G*^(*i,i*+1)^. *Then, for any loop γ in G*^(*i,i*+1)^ *based at u, the holonomy endomorphisms* 𝔽^(*i*)^(*γ*) *and* 𝔽^(*i*+1)^(*γ*) *are conjugate in* End_𝒞_ (𝔽(*u*)). *Consequently, the holonomy monoid conjugacy class (and any conjugacy-invariant statistic) is preserved across the overlap. The same holds for the fibre-jump indicator on overlap edges*.

*Proof*. Naturality implies *η*_*u*_ ° 𝔽^(*i*)^(*γ*) = 𝔽^(*i*+1)^(*γ*) ° *η*_*u*_ for the component *η*_*u*_ of the natural isomorphism at *u*, hence conjugacy. Fibre-jump invariance follows because *η* identifies fibres on overlap vertices and preserves isomorphism types.

#### Proposition 3.28

(Invariance under natural isomorphism / overlap comparison). *Let η* : 𝔽 ⇒ 𝔽^*′*^ *be a natural isomorphism of functors* Path(*G*_conf_) → 𝒞 . *Then for any loop γ* : *u* → *u*,

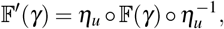

*so holonomy monoids are preserved up to conjugacy. Moreover, for any edge e* : *u* → *v*, 𝔽(*u*) ≅ 𝔽(*v*) *iff* 𝔽^*′*^(*u*) ≅ 𝔽^*′*^(*v*), *hence the fibre-jump indicator on edges is invariant under η*.

#### Proposition 3.29

(Holonomy conjugacy across overlap). *Let η* : 𝔽 | _*H*_ ⇒ 𝔽^*′*^ |_*H*_ *be a reconfiguration comparison on an overlap H. Then for any loop γ in H based at u*,

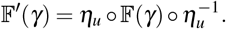

*Proof*. Naturality of *η* on each edge of *γ* implies commutation of *η* with the corresponding morphisms; composing around the loop yields the conjugacy relation.

#### Definition 3.30

(Rare structural reconfiguration criterion (ambient/interface coupling)). Fix at each stage a chosen ambient loop family ℒ_amb_ (loops supported away from *∂K*) and an interface loop family ℒ_*∂*_ (loops that visit the interface). A reconfiguration step is a *structural reconfiguration* if, after identifying the overlap via *η, both* the ambient holonomy diagnostic and the interface holonomy diagnostic change (e.g. the conjugacy class of *ρ*_*H*_ changes on representative loops from each family). This criterion encodes the principle that systemic reorganisation is rare and requires coupled changes in within-level organisation and boundary-mediated coupling.

#### Remark 21.

*In cyclic regimes such as a Hodgkin-type organisational cycle, one typically expects persistent holonomy without structural reconfiguration. Structural reconfiguration is reserved for rare events (e*.*g. long-term remodelling) and is introduced here primarily as a diagnostic notion*.

#### Remark 22

(Relation to geometric semantics). *Although we work in a combinatorial setting, the organisational bundle* 𝔽 *is a discrete analogue of bundle/connection structures used in geometric models of cortex and pattern formation [e*.*g. 53, 54]. The (scaffold, bundle) pair also admits a topos-theoretic reading in the “toposes as bridges” sense [5, 6], though we do not develop that embedding here*.

#### Operational outputs

Given a scaffold (*K, λ, B*), choose a derived base (*U, G*_conf_, supp, *p*) and a discrete bundle 𝔽 : Path(*G*_conf_) → 𝒞 . The framework yields mathematical outputs: (i) fibre-jump profiles and phase-space profiles along paths, (ii) loop types (stratum-pure/cross-level/interface), (iii) holonomy endomorphisms (monoid-valued in general) on selected loop families, and (iv) when a square-enriched 2-complex is specified and 𝒞 is additive, order-defect operators on chosen 2-cells. Along organisational histories, overlap comparisons transport these diagnostics and localise changes to explicit reconfiguration moves.

## 4 Hodgkin cycle as a multi-level organisational loop

We now instantiate the evolving organisational simplicial bundle framework of Section 3 on a single Hodgkin–Huxley spike. The aim is not to re-derive the classical Hodgkin–Huxley equations, but to provide a structurally explicit semantics for the multi-level pathway that Dennis Noble has emphasised as a paradigm of circular causality across levels and boundaries [43]. In particular we:

i. exhibit the spike as a *cross-level interface loop* in the configuration graph;
ii. identify a small number of *fibre jumps* along this loop, which encode changes of effective phase space (recruitment and coarse-graining of degrees of freedom);
iii. describe the corresponding *persistent holonomy* at a resting configuration and its biological interpretation;
iv. collect the resulting diagnostics in a case-study table (Table 5).

**Table 4:**
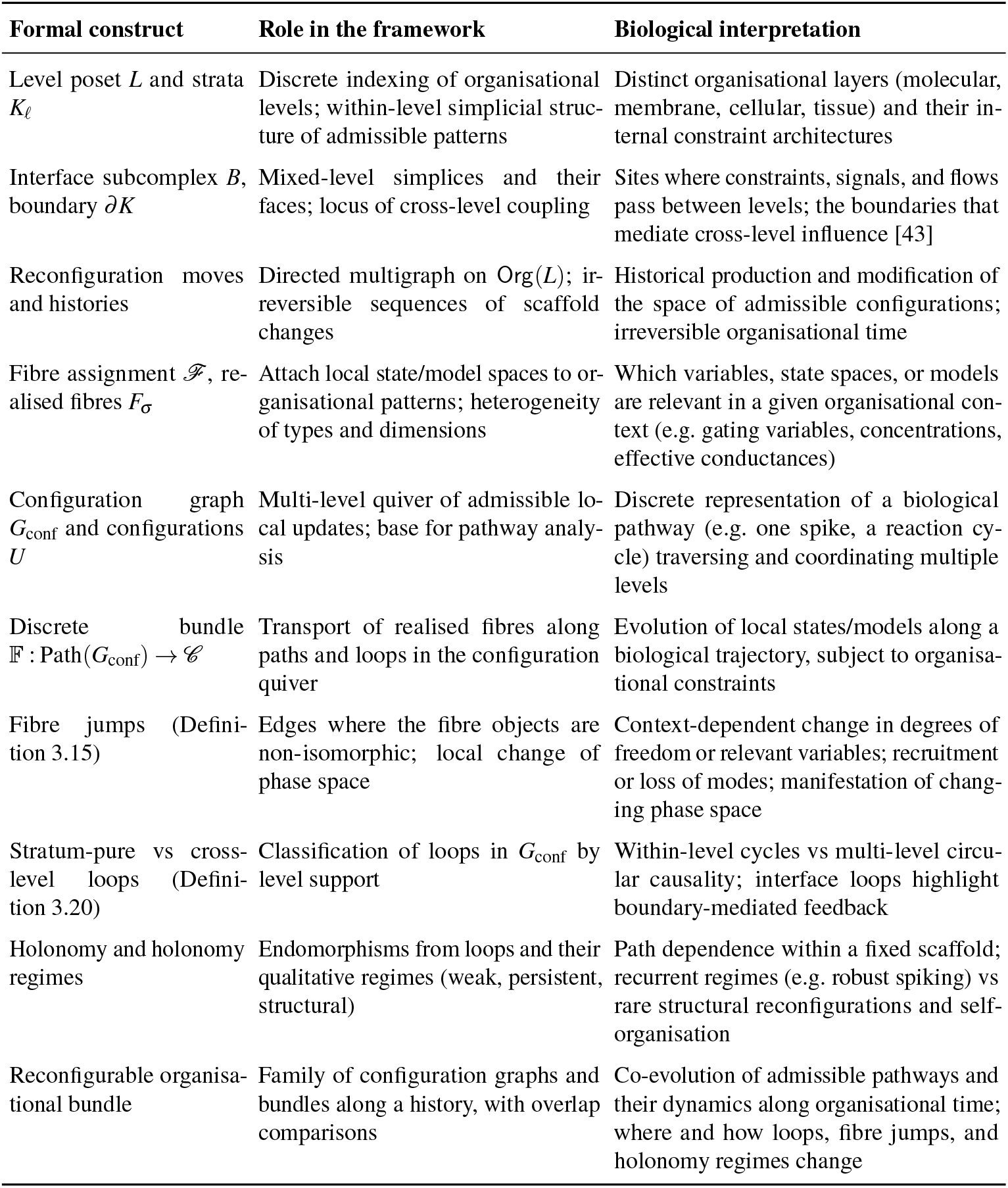
Crosswalk between the main formal constructs of evolving organisational simplicial bundles and their biological interpretation. Our aim is not to provide a complete modelling semantics for theoretical biology, but to give a first, mathematically explicit framework that takes key principles—changing phase space, non-entailing dynamics, multi-level constraints, and circular causality—as foundational [31, 33, 43].

**Table 5:**
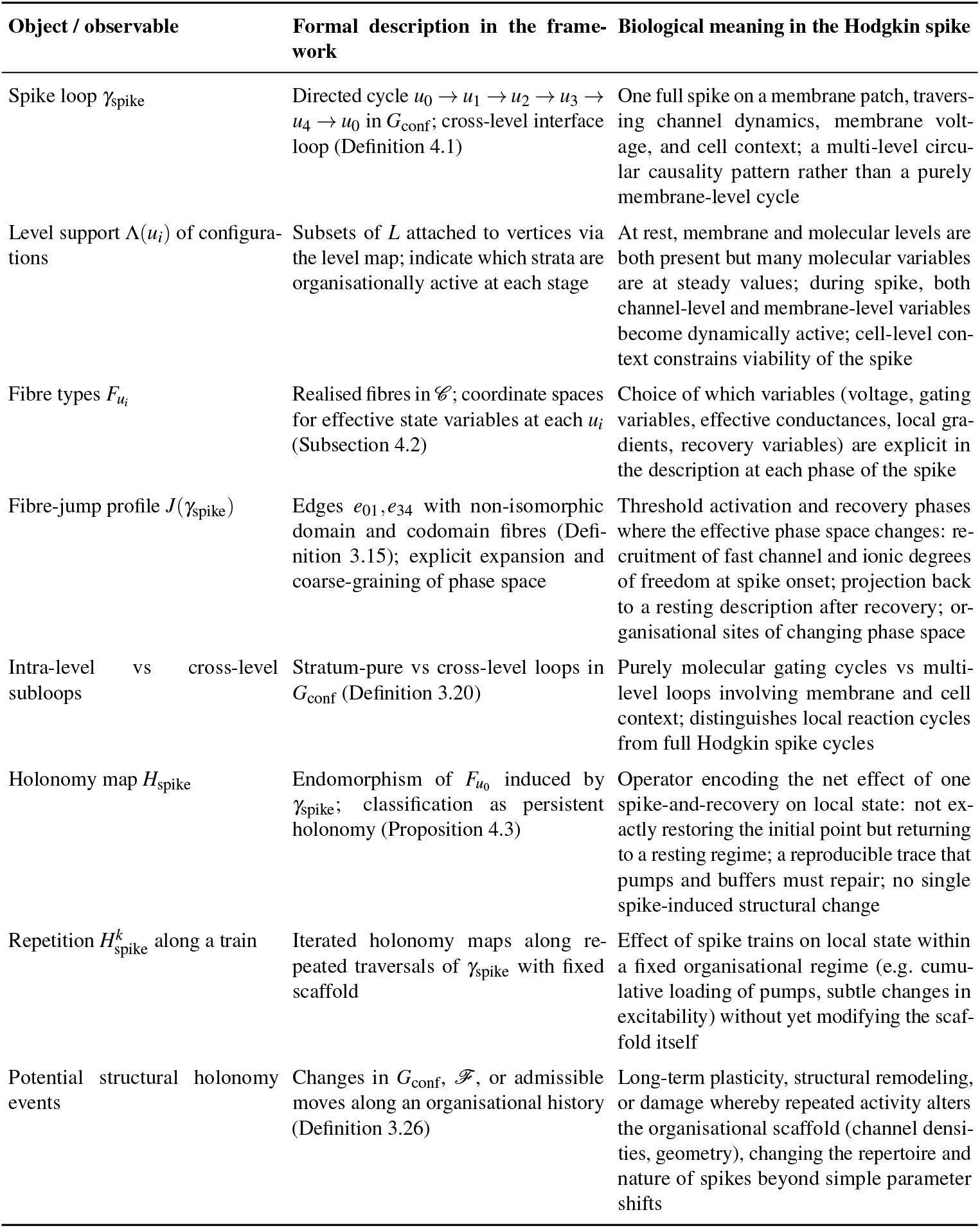
Case-study crosswalk between the formal constructs used in the Hodgkin section and their biological interpretation for a single spike. The spike appears as a cross-level interface loop with a non-trivial fibre-jump profile and persistent holonomy, providing a concrete realisation of multi-level circular causality and changing phase space in the sense of Longo and Montévil [31], Longo et al. [33], Noble [43].

This formalises Noble’s claim that the Hodgkin cycle is not a purely membrane-level cycle [44].

### Organisational setting for a single spike

We specialise the general scaffold to the classical Hodgkin picture on a small membrane patch of an excitable axon.

(A1) *Levels and strata*. We fix a level poset *L* = { *ℓ*_mol_ *< ℓ*_mem_ *< ℓ*_cell_ } corresponding respectively to molecular/channel complexes, membrane patch, and the local cell context. The organizational scaffold *K* then consists of:

- a membrane stratum 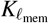 whose simplices encode local membrane regions and their adjacency;
- a molecular stratum 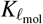 whose simplices encode channel complexes (Na^+^, K^+^, leak) and associated protein assemblies;
- a cellular stratum 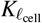 encoding the integration of the patch into the cell-wide environment (axonal cable, metabolic support).

The interface subcomplex *B* contains mixed simplices whose vertices lie in both 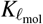 and 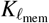, and in both 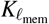 and 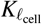, representing channel insertions into the membrane and coupling of the patch to the cell.

(A2) *Resting configuration*. We choose a reference organisational scaffold (*K, λ, B*) describing a patch in a stable excitable regime (classical Hodgkin–Huxley parameter range). This scaffold is held fixed for the analysis of a single spike: we do not consider plasticity or long-term structural change in this section.

(A3) *Configuration graph for one spike*. We restrict attention to a configuration graph (*U, G*_conf_, *p*) supported on:

- a resting configuration *u*_0_ (near-resting potential, typical channel distributions and states);
- an early upstroke / threshold configuration *u*_1_;
- a peak / early repolarisation configuration *u*_2_;
- a late repolarisation / recovery configuration *u*_3_;
- a recovered configuration *u*_4_ (restored sufficiently close to rest to support another spike).

The edges of *G*_conf_ include, in particular, a directed cycle

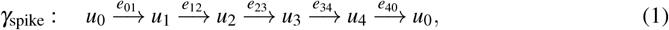

which we call the *spike loop*. Each configuration *u*_*i*_ is supported on a simplex *p*(*u*_*i*_) ∈ Simp(*K*), and its level support Λ(*u*_*i*_) = *λ* (*p*(*u*_*i*_)) ⊆ *L* typically involves both membrane and molecular levels, with *ℓ*_cell_ entering via the interface for current balance and metabolic context.

#### Definition 4.1

(Hodgkin spike as a cross-level interface loop). The configuration loop *γ*_spike_ in (1) is called the *Hodgkin spike loop* for the patch. Its level support 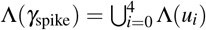 contains at least *ℓ*_mol_ and *ℓ*_mem_, and typically also *ℓ*_cell_; furthermore, several supporting simplices *p*(*u*_*i*_) intersect the interface boundary *∂K*. Thus, in the sense of Definition 3.20, *γ*_spike_ is a *cross-level interface loop*, not a stratum-pure loop.

This formalises Noble’s claim that the Hodgkin cycle is not a purely membrane-level cycle but a loop of mutual constraint across levels and boundaries [43].

### Fibre types, fibre jumps, and changing phase space

We now specialise the fibre assignment and identify a fibre-jump profile for the spike loop.

(B1) *Fibre category and realisation*. We fix the fibre category in this section to be 𝒞 = **FinVect**_ℝ_, the category of finite-dimensional real vector spaces and linear maps.

(B2) *Stage-wise fibre types*. For the spike loop *γ*_spike_, we consider the following fibre types at the configurations *u*_*i*_:

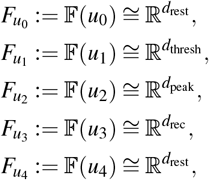

with each 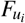 interpreted as follows:

- 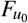 has coordinates for a resting potential, a small number of slow variables (e.g. effective leak conductance, slow channel population parameters), and summarised cell-context variables; many microscopic channel states and ion concentrations are not explicit because they are near steady.
- 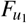 and 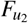 carry an expanded coordinate set, including fast Na^+^ and K^+^ gating variables and local ionic driving forces that become dynamically relevant during the upstroke and peak.
- 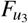 includes variables needed to describe recovery, e.g. K^+^ activation and Na^+^ inactivation states, plus a coarse representation of pump and buffering loads.
- 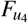 returns to the resting fibre type: fast variables are again close to a stable manifold and can be projected back into a smaller set of effective descriptors.

(B3) *Fibre-jump edges*. By Definition 3.15, an edge *e* : *u* → *v* of the configuration graph is a fibre jump if 𝔽(*u*) and 𝔽(*v*) are not isomorphic. In the spike loop we distinguish:

- *Threshold activation jump*. The edge *e*_01_ : *u*_0_ → *u*_1_ is a fibre jump: the effective phase space expands from 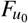 to 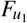 by making fast channel states and associated ionic driving forces explicit coordinates. Organisationally, the same patch now treats Na^+^ channels and the Na^+^ gradient as active degrees of freedom in the local dynamics.
- *Recovery/coarse-graining jump*. The edge *e*_34_ : *u*_3_ → *u*_4_ is also a fibre jump: fast degrees of freedom that were explicit in 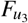 are projected back to an effective resting description 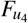 in which pumps and slower processes are summarised.

Along the other edges, *e*_12_ and *e*_23_, we model transport within a fixed phase-space type: we assume 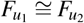 and 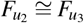, with the dynamics describing the upstroke and repolarisation in a common coordinate system.

#### Definition 4.2

(Fibre-jump profile of the spike loop). The *fibre-jump profile* of *γ*_spike_ is the ordered subset

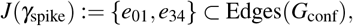

marking the threshold activation and recovery coarse-graining jumps.

Biologically, this profile displays where the effective phase space changes along the spike. The jump at *e*_01_ marks the onset of substantial dissipative ionic fluxes (Na^+^ influx, K^+^ efflux) at the level of the effective description; the jump at *e*_34_ marks where those fast processes become organisationally folded back into a smaller state space, with pumps and buffers restoring near-rest conditions. This refines the informal slogan of “changing phase space” in the spirit of Longo and Montévil [31], Longo et al. [33] by specifying *which* coordinates are added or removed at which steps.

### Holonomy of a single spike and its regime

Transport of fibre states along the configuration graph is encoded by the discrete organisational bundle 𝔽 : Path(*G*_conf_) → 𝒞 . For the spike loop, the associated holonomy map is

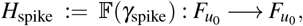

as in Definition 3.22.

We now make explicit the qualitative regime of this holonomy for a single spike in a healthy excitable cell.

#### Proposition 4.3

(Holonomy regime for a single Hodgkin spike). *Assume that:*

*(H1) the organisational scaffold* (*K, λ, B*) *and configuration graph* (*U, G*_conf_, *p*) *remain fixed during the spike (no plasticity or structural damage on the spike timescale);*

*(H2) the resting configuration u*_0_ *lies in a stable excitable regime (small perturbations in input produce spikes of comparable shape);*

*(H3) the bundle maps along the edges of γ*_spike_ *depend smoothly on a finite parameter set (channel densities, reversal potentials, temperature) and the spike loop persists under small parameter variations*.

*Then:*

i. *H*_spike_ *is not the identity: traversing the loop shifts the fibre state in* 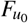 *by a small but non-zero amount, reflecting residual ionic and gating changes after a spike;*
ii. *H*_spike_ *is an endomorphism of* 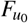 *in* 𝒞 *(monoid-valued holonomy): the spike-and-recovery loop maps a resting neighbourhood into a resting neighbourhood, but invertibility is not assumed at the level of coarse-grained transport;*
iii. *the qualitative spectrum and fixed-point structure of H*_spike_ *are invariant under sufficiently small parameter perturbations satisfying (H2)–(H3);*
iv. *consequently, the holonomy along γ*_spike_ *is* persistent *in the sense of Definition 3.25 and neither weak nor structural*.

#### Remark 23

(Biological interpretation of persistent holonomy). *In this case study, weak holonomy would mean that the spike leaves no macroscopically relevant trace at the level of the chosen coarse-graining, which is not physiologically accurate: a spike transiently alters ion distributions and channel states and engages pumps. Structural holonomy would require the spike to change the organisational scaffold itself (for example by modifying channel densities or membrane structure), which may occur under long-term activity or pathology but not for a single healthy spike. Persistent holonomy therefore encodes the intermediate situation: each spike enacts a reproducible but non-trivial transformation of local state, leaving the organisational scaffold and the repertoire of admissible pathways unchanged, yet producing a small “trace” that must be repaired by recovery mechanisms*.

#### Remark 24

(H_1_- and H_2_-type diagnostics for the spike loop). *On the subgraph H* ⊆ *G*_conf_ *whose vertices are the five spike configurations* {*u*_0_, …, *u*_4_} *and whose edges are those of γ*_spike_ *in* (1), *the underlying undirected graph is a single cycle. Topologically, the first homology group H*_1_(|*H*|; ℤ) *is isomorphic to* ℤ *and H*_2_( | *H* | ; ℤ) = 0. *Thus there is a single 1-dimensional generator corresponding to going once around the spike loop and no purely 2-dimensional obstruction in the base*.

*In the sense of the H*_1_*-type diagnostics of Definition 3.22, this reduces to the single holonomy element* 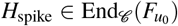 *the holonomy representation on the subgroup generated by γ*_spike_ *is completely determined (up to conjugacy) by H*_spike_. *Moreover, because our minimal spike graph does not include any commuting squares or alternative routes between configurations, the H*_2_*-type order-effect diagnostics are trivial on this restricted H: there are no specified* 2*-cells on which to measure non-commutativity. Non-trivial H*_2_*-type obstructions would appear only when the configuration graph is enriched with distinct pathways connecting the same pair of configurations (e*.*g. alternative molecular routes to repolarisation), which we do not model explicitly in this minimal single-spike case study*.

### Circular causality and internal time for the spike

The spike loop *γ*_spike_ is both a cross-level interface loop (Definition 4.1) and a holonomy-carrying loop (Proposition 4.3). In Noble’s terms [43], this expresses circular causality as follows:

- Membrane-level variables (voltage, local field) constrain which channel conformations and molecular interactions are active (interface from *ℓ*_mem_ to *ℓ*_mol_).
- Molecular-level channel dynamics and ion gradients feed back to membrane potential and current balance (interface from *ℓ*_mol_ to *ℓ*_mem_).
- The cell-level context (axonal cable, metabolic support) constrains viable spike shapes and recovery (interface between *ℓ*_mem_ and *ℓ*_cell_).

The cross-level nature of *γ*_spike_, the fibre-jump profile *J*(*γ*_spike_), and the persistent holonomy map *H*_spike_ together provide a mathematically explicit semantics for this multi-level circular causality.

From the perspective of internal time, a single spike sits at a fixed stage of an organisational history (Definition 3.6); the spike loop *γ*_spike_ generates one instance of the *circular axis* of time (Remark 20) via repeated traversals. In a train of spikes with a fixed scaffold, the internal time evolution is described by the powers 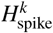 acting on resting states, while slow plasticity or pathology would correspond to occasional reconfiguration moves that alter the configuration graph, the fibre assignment, or the holonomy regime.

In the Hodkgin example, the two components of internal biological time from Remark 20 become concrete. Along a fixed organisational stage, with scaffold (*K, λ, B*) and bundle 𝔽 held fixed, the spike loop *γ*_spike_ generates a circular time axis: iterating the spike corresponds to iterating the holonomy map *H*_spike_ on the rest fibre, and hence to repeated traversals of the lifted trajectory in *E*^spike^. Along an organisational history of reconfigurations, the scaffold and bundle data change, so that the configuration graph, the spike loop, and the associated holonomy *H*_spike_ slowly deform. The irreversible axis of organisational histories thus records how the repertoire and geometric shape of spiking loops evolve, while the circular axis records the action of individual spikes on local state spaces. The interaction of these two axes realises, in a precise combinatorial–geometric form, the theoretical-biology view in which rhythms and cycles (spikes) are nested within an irreversible history of changing phase spaces and organisational constraints [31, 43].

### Geometry of the lifted spike loop in the total space

For the purpose of this subsection, it is convenient to make explicit the “total space” associated with a discrete organisational bundle over the configuration graph. Given a configuration graph (*U, G*_conf_, *p*) and a discrete bundle *F* : Path(*G*_conf_) → 𝒞 as in Definition 3.13, we can form the set

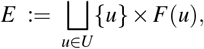

with the obvious projection *π* : *E* → *U, π*(*u, x*) = *u*. We refer to *E* as the (discrete) total space of the organisational bundle.

Restricting to the Hodgkin spike loop, we obtain the finite sub-bundle

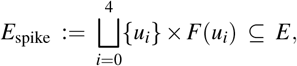

with base the five spike configurations *u*_0_, …, *u*_4_ and fibres *F*(*u*_*i*_) as in §4. A *lifted spike trajectory* is a path in *E*_spike_ of the form

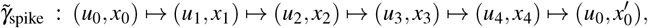

such that each step projects to the corresponding edge of *γ*_spike_ and (*u*_*i*+1_, *x*_*i*+1_) is obtained from (*u*_*i*_, *x*_*i*_) by the bundle map *F*(*e*_*i,i*+1_).

On the segments of *γ*_spike_ where there is no fibre jump (*e*_12_ and *e*_23_ in our minimal model), the fibres *F*(*u*_*i*_) are isomorphic and the lifted trajectory moves within a fixed finite-dimensional space (upstroke and repolarisation in a common coordinate system). At the threshold activation and recovery edges *e*_01_ and *e*_34_, the lifted trajectory crosses between non-isomorphic fibres: geometrically, 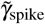 passes through regions of *E*_spike_ where the effective number or nature of coordinates changes. These steps correspond to the recruitment and subsequent coarse-graining of fast channel and ionic degrees of freedom discussed in §4.

The holonomy operator *H*_spike_ = *F*(*γ*_spike_) can be viewed as the net effect of one turn of 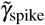 on the resting fibre *F*(*u*_0_). Even though the base loop *γ*_spike_ is combinatorially simple, the lifted trajectory in *E*_spike_ is genuinely stratified: it is a concatenation of segments in different fibres glued by the jump maps at *e*_01_ and *e*_34_. In this sense, the spike is *not* merely a closed orbit in a fixed phase portrait, but a loop in the total space that traverses different local phase spaces in a coherent way.

From the viewpoint of circular causality, this total-space picture makes explicit that the spike loop combines:

- motion within a given organisational level and fibre (e.g. evolution of gating variables and voltage during upstroke and repolarisation), and
- discrete changes of organisational context and phase space encoded by fibre jumps at cross-level interfaces.

Biologically, the resulting loop in *E*_spike_ captures, in a single object, how membrane voltage, channel conformations, ionic gradients and cell context are jointly reorganised over the course of one spike, providing a geometric counterpart to Noble’s description of the Hodgkin cycle as a multi-level loop rather than a mere membrane-level cycle.

### Homological diagnostics for the spike loop

At the level of the configuration graph, the single-spike loop *γ*_spike_ defines a homology class

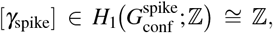

which can be taken as a generator of the first homology group of the spike subgraph. The existence and stability of such a generator are therefore a first diagnostic of a robust spiking regime: under admissible reconfigurations of the scaffold and the induced configuration graph, the persistence of a non-trivial class [*γ*_spike_] signals that the basic organisational loop is preserved.

At the level of the organisational scaffold, the spike loop projects to a family of loops in the stratified simplicial complex (*K, λ, B*), each encircling part of the interface boundary *∂K*. Non-trivial classes in *H*_1_(*K*; ℤ) that are represented by such projections provide a complementary diagnostic: they detect recurrent organisational patterns in the scaffold itself, rather than in a particular pathway.

Higher homology groups become relevant when one considers families of spikes or extended patterns. For example, if a two-parameter family of spikes fills in a surface in the configuration graph, then obstructions to contracting this surface, captured by non-trivial elements of *H*_2_(*G*_conf_; ℤ), indicate that the spike loop is constrained by a higher-order organisational structure (for instance, a metabolic or electrophysiological manifold of admissible regimes). In the present paper we restrict attention to the basic generator [*γ*_spike_] ∈ *H*_1_, but the framework supports such higher-dimensional diagnostics in principle.

### Total-space view of a single spike

Restrict attention to the configurations visited by the spike loop *γ*_spike_. Let

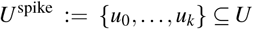

and let 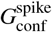 be the subgraph of *G*_conf_ induced by *U* ^spike^. The corresponding *total space for the spike* is the restricted bundle

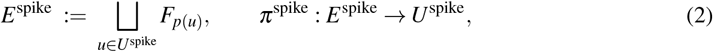

together with the transport maps 𝔽(*e*) for edges *e* in 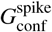.

It is convenient to organise *E*^spike^ into slices according to the level support. Writing *L*_chan_, *L*_mem_, *L*_cell_ ∈ *L* for the channel, membrane, and cellular levels, we set

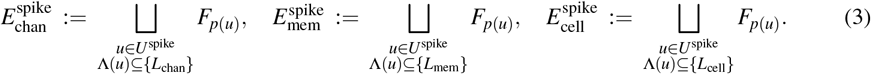

Over configurations whose support intersects several levels, the fibres are attached along interface simplices in ∂*K*; in this sense *E*^spike^ is obtained by gluing level-wise slices along mixed-level fibres over the interface boundary.

The lifted spike trajectory is then a path in *E*^spike^ that alternates between within-level segments and cross-level ramps. Starting from a rest state 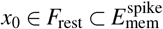 with *π*^spike^(*x*_0_) = *u*_0_, we obtain a sequence

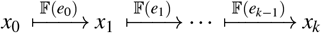

with *π*^spike^(*x*_*i*_) = *u*_*i*_ and *x*_*k*_ = *H*_spike_(*x*_0_) as in (4). Segments corresponding to stratum-pure edges (Definition 3.20) move within a single slice 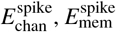 or 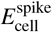, whereas segments corresponding to interface edges lift to ramps that connect slices over the interface boundary ∂*K*.

This picture makes precise why the Hodgkin spike is not merely a loop in a fixed phase space: the loop *γ*_spike_ in base space lifts to a zigzag trajectory in the stratified total space *E*^spike^ that repeatedly passes through gluing zones where the fibre type changes. The net effect of the spike on the rest fibre is the holonomy *H*^spike^, rather than a return to the identical microscopic state. In this sense the total-space view provides a mathematical semantics for Noble’s multi-level circular causality through boundary-mediated feedback [33, 43].

### Holonomy of a single spike

Let 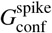 denote the subgraph of the configuration graph generated by the vertices and edges of the single-spike loop *γ*_spike_ : *u*_0_ → *u*_1_ → … → *u*_*k*_ = *u*_0_. We choose *u*_0_ to be a rest configuration and write *σ*_rest_ := *p*(*u*_0_) ∈ Simp(*K*) for its distinguished supporting simplex, with realised fibre

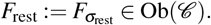

The discrete organisational bundle 𝔽 : Path(*G*_conf_) → 𝒞 of Definition 3.13 restricts to a functor on 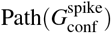, which we still denote by 𝔽. The *holonomy map* of a single Hodgkin spike is the endomorphism

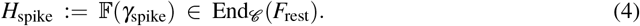

By construction, a rest state *x*_0_ ∈ *F*_rest_ is transported along the lifted spike trajectory to a final state

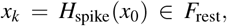

which in general need not coincide with *x*_0_.

#### Remark 25

(Holonomy regime for the Hodgkin spike). *In the organisational regimes of interest, the rest fibre F*_rest_ *encodes the effective parameters of an excitable membrane segment at rest (for example, coarse-grained conductances and inactivation variables). A single spike produces a small but reproducible update of these parameters (refractoriness, partial inactivation) without destroying the excitable regime. In our framework this is expressed by assuming that:*

i. *H*_spike_ *is not the identity on F*_rest_, *i*.*e. the spike leaves a non-trivial trace on the rest state; and*
ii. *the qualitative type of H*_spike_ *(e*.*g. its fixed-point structure or spectrum when* 𝒞 *is linear) is stable under admissible perturbations of the bundle data*.

*Under these assumptions the single-spike loop γ*_spike_ *realises a regime of* persistent holonomy *in the sense of Definition 3.25. Weak holonomy would correspond to a spike leaving the rest state exactly unchanged, whereas a transition to structural holonomy would signal a rare reconfiguration of the underlying organisational scaffold (for example, long-term remodelling or pathology) rather than the dynamics of an individual spike*.

## 5 Discussion and outlook

We introduced a discrete, combinatorial framework of *evolving organisational simplicial bundles* and instantiated it on a single Hodgkin spike. Table 6 summarises how these constructs align with Noble’s principles and with modelling desiderata from theoretical biology.

**Table 6:**
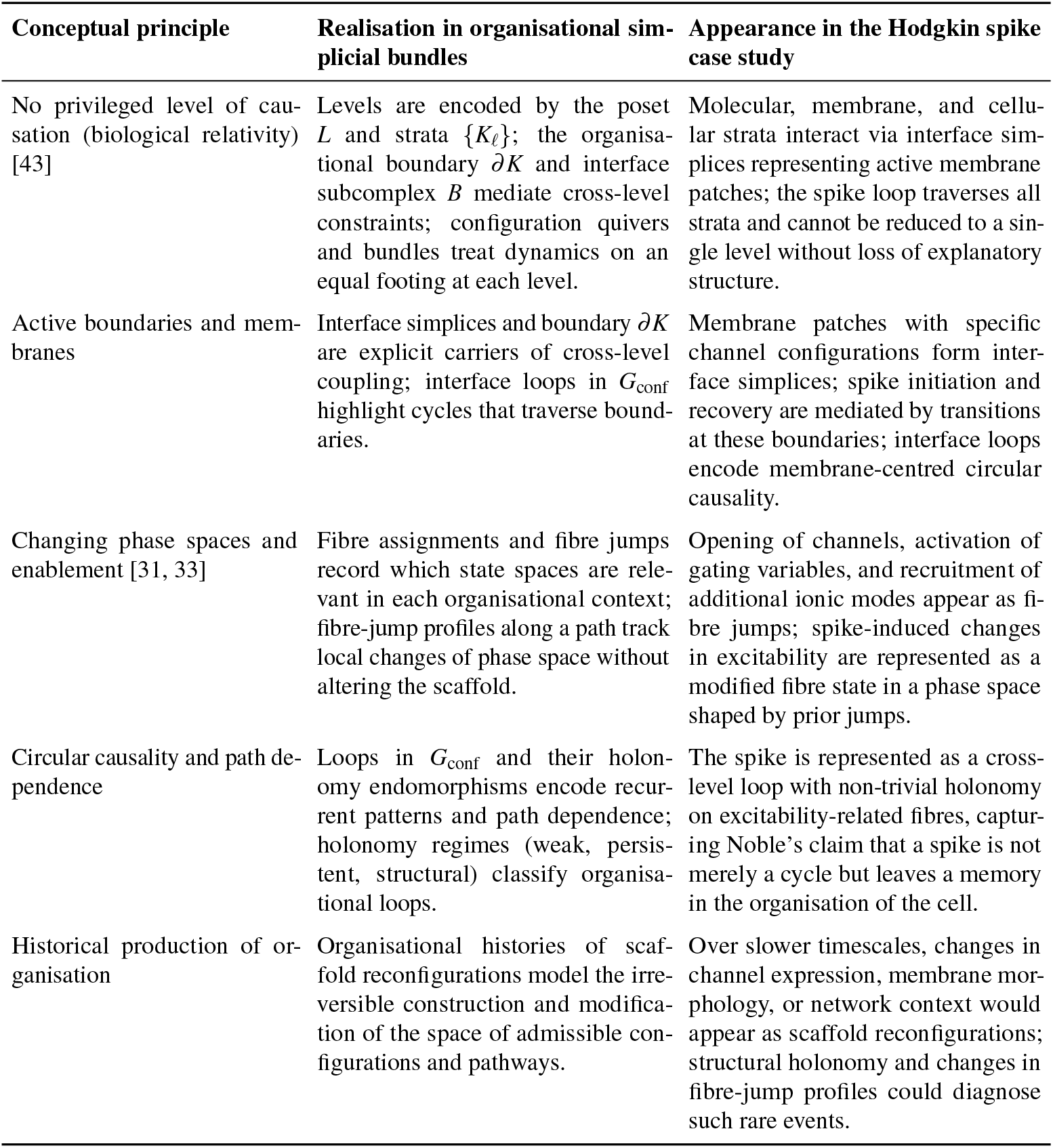
Alignment between key conceptual principles from theoretical biology and the organisational simplicial-bundle framework, with their realisation in the Hodgkin spike case study.

### Limitations and further work

The present paper is intentionally a first semantics step, and its main limitation is the restricted treatment of biological heterogeneity—both in the *organisational space* and in the *state descriptions* carried by fibres.

#### Heterogeneous organisational space

We have modelled the organisational scaffold by a single simplicial complex with a stratification and interface data. In broader biological settings, local regions may admit such descriptions while the global organisation is far more irregular: the effective “geometry” can be non-manifold, highly nonuniform across scales, and not representable by a single globally coherent combinatorial type. A natural next step is to treat organisational contexts as forming a *site* and to represent organisation by descent/gluing data over that site, so that the admissible local models and their overlaps are part of the semantics rather than a fixed background choice.

#### Heterogeneous fibres and changing data types

We have also fixed a single target category 𝒞 for fibres. Real systems demand stronger heterogeneity: fibre types may vary across strata and, crucially, along histories (e.g. a region whose effective state description is a set or finite-state object at one stage may require probabilistic simplices, constraint manifolds, or other structured objects at another). Capturing such *type change* suggests replacing a single 𝒞 by an indexed/categorical semantics varying with context— for example via fibrations or, more systematically, by moving to a Grothendieck-topos setting where objects, morphisms, and internal logic are context-sensitive as in [3]. This is precisely the direction in which “toposes as bridges” methodologies become relevant: they provide a disciplined way to internalise heterogeneity and to relate multiple modelling languages within a common semantic framework [5, 6].

Beyond heterogeneity, three further developments are immediate. First, we have treated 𝔽 as given; incorporating stochastic or deterministic dynamics on fibres should be done by *constraining* families of mechanistic models with organisational semantics, rather than replacing them. Second, our cohomological diagnostics are not operational; a systematic development (including stability under refinement/coarse-graining) remains open. Third, the Hodgkin example is deliberately minimal; extending to multi-spike patterns, plasticity, and network/tissue organisation will provide sharper tests of fibre-jump and holonomy diagnostics.

Finally, the framework is directly compatible with several ongoing programmes: autonomy and minimal living systems emphasising membranes and closure of constraints [49, 50], geometric approaches to neural organisation developed by Sarti and collaborators [53, 54], and (as argued above) topos-theoretic unification as the appropriate next step for representing biological heterogeneity without collapsing it to a single homogeneous state space [5, 6] and biological relativity [42, 43].

## Acknowledgements

The author thanks Giuseppe Longo for his comments and discussions during the course of this work.

## A Supplementary Material

### Diagnostic pipeline

For the Hodgkin cycle it is useful to summarise how the foregoing structures are used operationally. Given a realised organisational history and a chosen pathway, the diagnostic pipeline is:

(D1) Fix a scaffold snapshot (*K, λ, B*) and construct a configuration graph (*U, G*_conf_, supp, *p*) satisfying the axioms of Definition 3.9.

(D2) Choose a fibre assignment ℱ and realisation *R* into 𝒞 = **FinVect**_ℝ_, obtaining realised fibres *F*_*σ*_ and a discrete bundle 𝔽 : Path(*G*_conf_) → 𝒞.

(D3) Compute the fibre-jump profile (Definition 3.15) and basic invariants such as the dimension profile along selected paths and loops. This identifies where the local phase space changes along the pathway.

(D4) Analyse loop structure in *G*_conf_ (Definition 3.20), distinguishing stratum-pure, cross-level, and interface loops, and compute their holonomy images under 𝔽 (Definition 3.22). Use the H^1^-type diagnostics to compare different trivialisation choices, and H^2^-type order-defect diagnostics to localise order effects.

(D5) Along an organisational history, track how configuration graphs, fibre-jump profiles, and holonomy regimes (Definition 3.25) change under reconfiguration of the scaffold (Definition 3.26). Rare transitions to new holonomy regimes or qualitative changes in fibre-jump structure are interpreted as candidate signatures of structural change or self-organisation.

### Summary table: formal constructs and biological interpretation

Table 4 summarises the main constructs introduced in this section and their biological interpretation.

### Case-study diagnostics and biological interpretation

Table 5 summarises the main formal constructs and observables for the Hodgkin case study and their biological interpretation.

